# Serine 1283 in extracellular matrix glycoprotein Reelin is crucial for Reelin’s function in brain development

**DOI:** 10.1101/2022.01.06.474241

**Authors:** Ralf Kleene, Gabriele Loers, Ahmed Sharaf, Shaobo Wang, Hardeep Kataria, Max Anstötz, Irm Hermans-Borgmeyer, Melitta Schachner

## Abstract

Deficiency in the extracellular matrix glycoprotein Reelin severely affects migration of neurons during development. The function of serine at position 1283 in Reelin has remained uncertain. To explore its relevance we generated *rln*^*A/A*^ mice that carry alanine instead of serine at position 1283, thereby disrupting the putative casein kinase 2 (CK2) phosphorylation site S_1283_DGD. Mutated mice displayed *reeler*-like locomotor behavior, abnormal brain anatomy and decrease of Reelin RNA and protein levels during development and in adulthood. Since serine 1283 was previously proposed to mediate proteolysis of adhesion molecules, we investigated proteolysis of cell adhesion molecule L1 and found it normal in *rln*^*A/A*^ mice. Neuronal migration in the embryonic *rln*^*A/A*^ cerebral cortex was impaired, but rescued by *in utero* electroporation of the Reelin fragment N-R6 containing the putative CK2 phosphorylation site. In *rln*^A/A^ mice migration of cerebellar granule cells *in vitro* was promoted by application of wild-type but not by mutated Reelin. In cerebellar neuron cultures, Reelin expression was decreased upon inhibition of ecto-phosphorylation by CK2. Biochemically purified wild-type, but not mutated Reelin was found phosphorylated. Altogether, the results indicate that ecto-phosphorylation at serine 1283 rather than proteolytic processing of adhesion molecules by Reelin plays an important role in Reelin functions.

## 1. Introduction

The laminated architecture of brain structures, such as the cerebral cortex and the cerebellum, results from the orchestrated migration of neurons. The extracellular matrix protein Reelin controls the sequential lamination of the cerebral cortex by regulating the migration of newborn neurons [1]. Reelin is also important for the postnatal lamination and foliation of the cerebellum which is important for movement, cognition, and language [2]. The naturally arisen Reelin-deficient mouse mutant *reeler* shows disorganized cortical layers, severe cerebellar malformation with hypoplasia, ataxia, loss of balance, and reeling gait [3-8]. Reelin promotes neuronal differentiation, synapse formation and synaptic plasticity, and malfunction of Reelin was found to be associated with multiple neurological problems and behavioral disorders in humans [9-13].

Due to the functional cooperation between different proteases, Reelin is specifically cleaved at the N-terminus within Reelin repeat 3 and at the C-terminus between Reelin repeats 6 and 7 [14-17]. This processing leads to the generation of five different Reelin fragments that regulate the duration and range of Reelin functions, e.g. in adhesion and in de-adhesion [14, 16, 18-22]. It has been reported that Reelin shows serine protease activity and can cleave adhesion molecules [23-25]. The serine at position 1283 in the third Reelin repeat was proposed to be the catalytic center [26] and the exchange of serine 1283 to alanine disrupts the proteolytic cleavage of the cell adhesion molecule L1 *in vitro* [25]. Since the Reelin sequence S_1283_DGD represents an S/T-D/E-X-E/D motif for phosphorylation by ecto-casein kinase 2 (CK2) [27], it was conceivable that phosphorylation of serine 1283 regulates Reelin’
ss functions and that disruption of this phosphorylation site impairs its functions. To study the relevance of serine 1283 *in vivo*, we generated a gene-edited mouse with an exchange of serine 1283 against alanine. We also performed *in vitro* experiments with cerebellar granule cells to allow for an understanding of the molecular mechanisms underlying the functions of serine 1283. We show the importance of Reelin fragment N-R6, which comprises the central fragment and the N-terminal fragment N-R2 and contains the putative casein kinase 2 phosphorylation site at serine 1283.

## 2. Results

### 2.1. Disruption of serine 1283 in Reelin results in brain malformation and behavioral abnormalities in gene-edited mice

We used the Zinc Finger Nuclease technique to generate gene-edited mice expressing Reelin with the S/A_1283_ mutation (Fig. 1a). Insertion of the donor DNA carrying the mutated Reelin sequence into the Reelin gene introduced a restriction site for AciI. Mice homozygous for the mutation (*rln*^*A/A*^) were identified by genotyping using genomic DNA for PCR and analyzing the PCR products after treatment with the restriction enzyme AciI. Wild-type (*rln*^*+/+*^) mice that were derived from the breeding of heterozygous (*rln*^*+/A*^) mice showed only the undigested 530 bp amplicon, while the homozygous mutation-carrying *rln*^*A/A*^ mice showed the 199 bp and 331 bp AciI digestion products (Fig. 1b). In *rln*^*+/A*^ mice, both the 530 bp amplicon and the 199 bp and 331 bp fragments were observed (Fig. 1b). Whole genome sequencing and Southern blot analysis (Fig. 1c) showed that the donor DNA was correctly inserted into the Reelin gene, and no additional insertions were detected. Gene sequencing also showed correct insertion of the donor DNA into the Reelin gene (Supplementary Figure 1). At postnatal day 18 *rln*^*A/A*^ mice were reduced in body size and weight by approximately 40% when compared with *rln*^*+/+*^ littermates (Fig. 1d, e). In addition, *rln*^*A/A*^ mice showed severe deficits in motor functions, including tremor (Supplementary Movie 1). Other behavioral abnormalities were a general lack of coordination, including reduced upright stance and explorative drive (Fig. 1f; Supplementary Movie 1), disorientation and inability to overcome hurdles, in comparison to their normal appearing *rln*^*+/A*^ and *rln*^*+/+*^ littermates (Fig. 1f; Supplementary Movie 1). In the pencil-grab test, *rln*^*A/A*^ mice failed to grab the pencil with their forepaws, and their hind limbs were stiff with pronounced discoordination (Fig. 1g).

**Figure 1.**
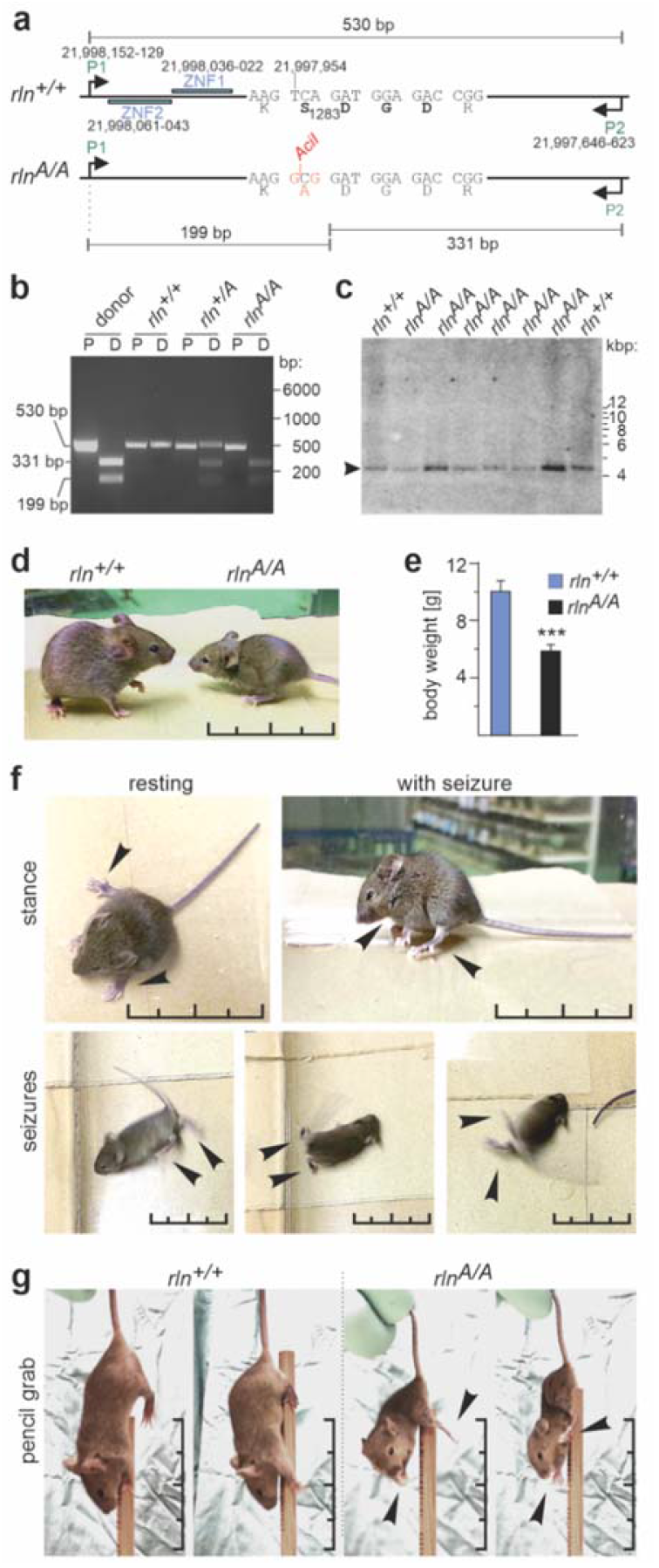
Generation of mice expressing Reelin with the S/A_1283_ mutation. (**a**) Targeted mutation and genotyping strategies: the wild-type *Reelin* gene sequence (*rln*^*+/+*^) was used to design zinc finger nucleases 1 and 2 (ZFN1 and ZFN2) which bind close to the codon of serine 1283 (TCA coding for serine). Genomic positions are indicated. In the mutated sequence (*rln*^*A/A*^) TCA is replaced by GCG (coding for alanine), thereby generating a restriction site for AciI. Primers P1 and P2 were used to amplify a 530 bp region as indicated. After digestion with AciI, the 530 bp amplicon was cleaved to yield the 199 bp and 331 bp fragments. (**b**) Genotyping of gene-edited mice; P: amplicons with primers P1 and P2; D: AciI-digested amplicon; the pUC vector carrying the donor sequence served as control (donor). (**c**) Southern blot analysis of gene-edited mice; only the 4135 bp EcoRI fragment of the Reelin gene (arrowhead) was detected in two *rln*^*+/+*^ mice and six *rln*^*A/A*^ mice using a radiolabeled 530 bp probe derived from PCR amplification using primers P1 and P2 and the pUC vector carrying the donor sequence. See also Supplementary Figure 1. (**d**-**g**) Phenotype of one-month-old mice expressing Reelin with the S/A_1283_ mutation: *rln*^*A/A*^ mice display reduced body size (**d**) and body weight (**e**). Mean value + SEM are shown for the body weight (***p<0.0001; Student’s t-test) (**e**). *rln*^*A/A*^ mice are deficient in upright stance and motor coordination with frequent episodes of tremor (**f**). *rln*^*A/A*^ mice fail to grab a pencil with their fore- and hindpaws (**g**). (**f, g**) Arrowheads indicate regions of impairment. (**d, f, g**) Scale bars: 5 cm. (**d**-**g**) 14 animals per genotype were used. See also Supplementary Movie 1.

### 2.2. Disrupted layering in cerebral cortex, hippocampus and cerebellum of rln^A/A^ mice

In comparison to the brains from *rln*^*+/+*^ mice, the gross anatomy of *rln*^*+/A*^ brains was normal, whereas *rln*^*A/A*^ brains showed an abnormally developed mesencephalon and cerebellum (Fig. 2a). The mesencephalic superior and inferior colliculi as well as cisternae ambiens and magna were enlarged, and the cerebellum was reduced in size and lacked the characteristic foliation and fissures (Fig. 2a). Nissl staining revealed a normally developed cortex and hippocampus in *rln*^*+/+*^ mice, while *rln*^*A/A*^ mice showed a distorted cytoarchitecture of the hippocampus with scattered granule cells and malpositioned pyramidal neurons and a *reeler*-like cell-rich area in the area of the cortex that is analogous to the marginal zone in the *rln*^*+/+*^ cerebral cortex (Fig. 2b). Layer-specific immunostaining for upper cortical layers revealed *reeler*-like displacement of Cux1- and Brn2-positive cells (Fig. 2c, d). To investigate expression of mutated Reelin in Ca-jal-Retzius cells, *rln*^*A/A*^ mice were cross-bred with *CXCR4-GFP* mice that express GFP specifically in Cajal-Retzius cells [28, 29], and sections from the cerebral cortex of these mice were analyzed.

**Figure 2.**
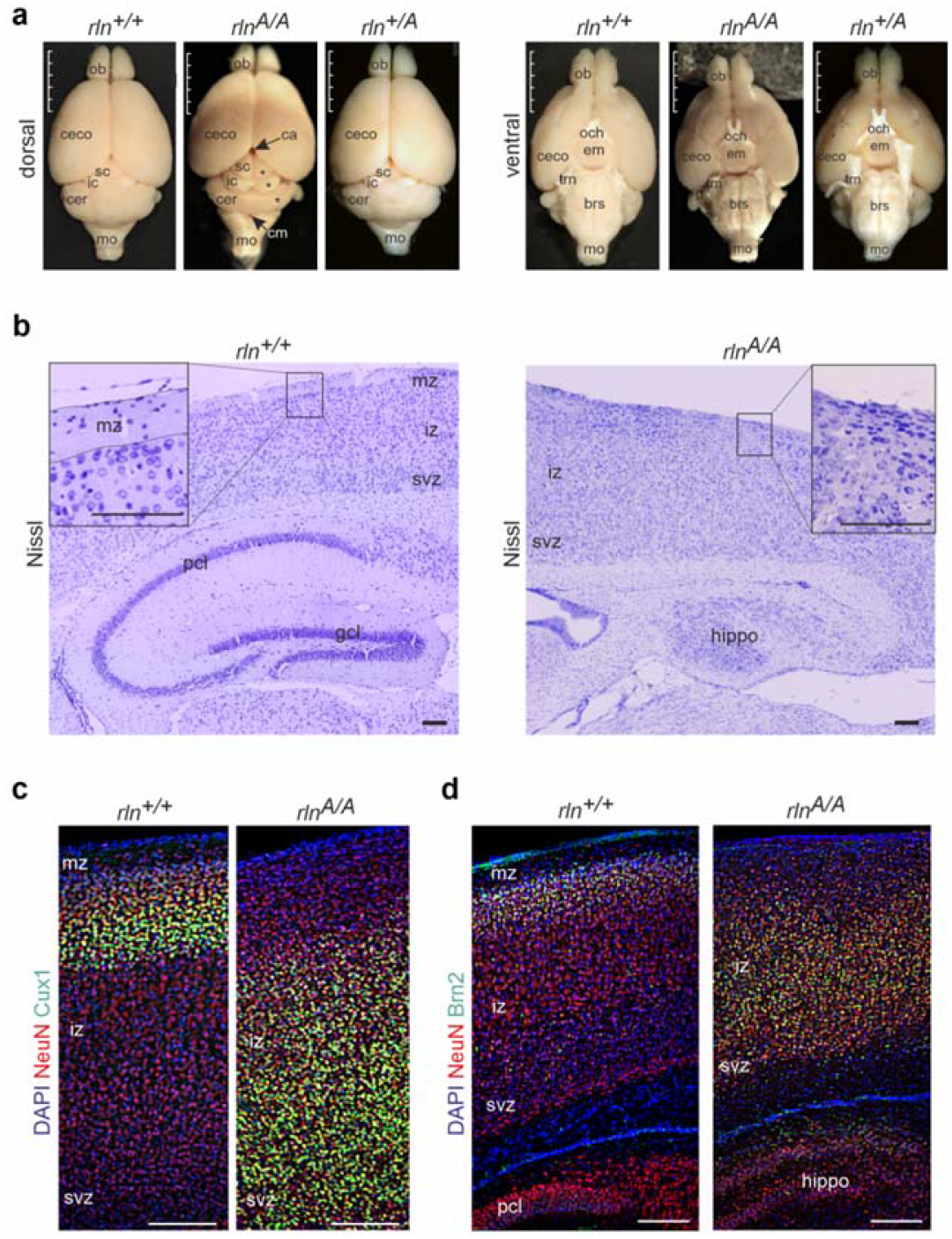
Abnormal shape of brains and cytoarchitecture of the cerebral cortex and hippocampus in *rln*^*A/A*^ mice. (**a**) Appearance of brains from one-month-old *rln*^*A/A*^ mice shows an enlarged superior colliculus (sc) and inferior colliculus (ic), enlarged cisterna ambiens and magna, and a smaller and unfoliated cerebellum (cer) when compared to normal brains from *rln*^*+/+*^ and *rln*^*+/A*^ littermates. Scale bars: 5 mm. Asterisks indicate the abnormal mesencephalon and cerebellum. Arrows indicate the enlarged cisterna ambiens (ca) and cisterna magna (cm). brs - brain stem; ceco - cerebral cortex; em - eminentia mediana; ob - olfactory bulb; mo - medulla oblongata; och - optic chiasm; trn - trigeminal nerve. (**b**) Nissl staining indicates a distorted layer organization of the cerebral cortex and hippocampus of *rln*^*A/A*^ mice when compared to *rln*^*+/+*^littermates. A region in the *rln*^*A/A*^ cortex analogous to the marginal zone (mz) of the *rln*^*+/+*^ cortex is densely populated by neurons. The regions in the *rln*^*A/A*^ hippocampus (hippo) which correspond to the granule cell layer (gcl) and pyramidal cell layer (pcl) of the *rln*^*+/+*^ hippocampus contain dispersed granule and pyramidal cells; iz, intermediate zone; svz, subventricular zone; scale bars: 100 μm. The border between svz and iz is indicated by bright field light microscopy as a distinct border between the white matter of the corpus callosum and the grey matter of the cerebral cortex. (**c, d**) Layer-specific immunostaining for the upper layers of the cerebral cortex with antibodies against Cux1 or Brn2 (green) in combination with immunostaining of neurons using NeuN (red) antibody reveals abnormally laminated *rln*^*A/A*^ cerebral cortices in comparison to normally laminated cerebral cortices of *rln*^*+/+*^ littermates; nuclear staining with DAPI (blue); scale bars: 100 μm. (**a**-**d**) Four animals per genotype were used.

In *rln*^*A/A*^-*CXCR4-GFP* cerebral cortices, most of the Cajal-Retzius cells were positioned closer to the pial surface than the Cajal-Retzius cells in *rln*^*+/+*^*-CXCR4-GFP* controls and some of the *rln*^*A/A*^ Cajal-Retzius cells were misplaced and in lower layer positions (Fig. 3a, b). Not only the cerebral cortex, but also the cerebellum of *rln*^*A/A*^ animals appeared severely hypoplastic and was *reeler*-like with disorganized morphology (Fig. 3c) with misplaced calbindin-positive Purkinje cells and dispersed cerebellar granule cells (Fig. 3d; Supplementary Figure 2). These results suggest that mutation of serine at 1283 in Reelin entails a *reeler*-like phenotype in mice.

**Figure 3.**
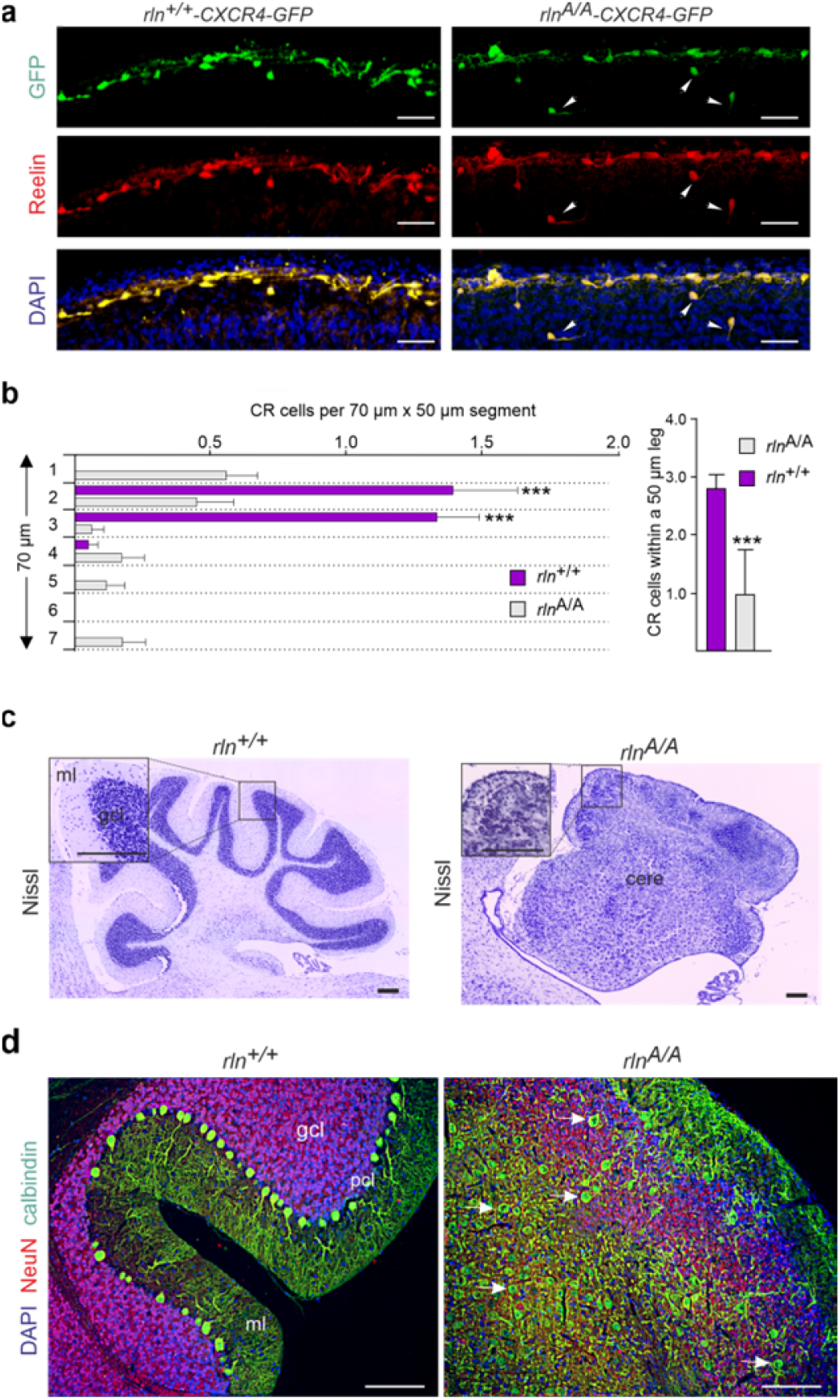
Misplacement of Cajal-Retzius and Purkinje cells *rln*^*A/A*^-*CXCR4-GFP* mice. (**a**) Immunostaining with Reelin antibody G10 shows Reelin expression (red) in Cajal-Retzius cells (green) in the marginal zone of the cerebral cortex of one-month-old *rln*^*+/+*^-*CXCR4-GFP* mice and in the analogous region in one-month-old *rln*^*A/A*^-*CXCR4-GFP* mice. Several partially displaced Ca-jal-Retzius cells (white arrows) were observed in the *rln*^*A/A*^*-CXCR4-GFP* mice in the marginal zone (mz); scale bars: 30 μm. (**b**) Numbers of Cajal-Retzius cells in the marginal zone of the somatosensory cortex in *rln*^*+/+*^*-CXCR4-GFP* mice and in the corresponding analogous region of *rln*^*A/A*^*-CXCR4-GFP* mice. The marginal zone in *rln*^*+/+*^ mice and the analogous area in *rln*^*A/A*^ mice was divided into 4-5 segments and each segment was further divided into 7 bins. Cajal-Retzius cells in each bin were counted and their distribution across the 7 bins of each segment was assessed (left panel). In addition, the numbers of Cajal-Retzius cells were counted in at least 5 legs of 50 μm in the mz of *rln*^*+/+*^ mice and in the analogous area of *rln*^*A/A*^ mice (right panel). Mean values + SEM are shown for the number of Cajal-Retzius cells in a 50 μm leg (n=18; ***p<0.0001; two-tailed Student’s t-test) and in a 70×50 μm segment (n=18; ***p<0.0001; two-way ANOVA for multiple comparisons and Sidak’s multiple comparisons test). (**c, d**) Nissl stain and immunostaining for calbindin (green) and NeuN (red) reveals severe malformations in the *rln*^*A/A*^ cerebellum (cere): lack of a distinct Purkinje cell layer (pcl) and granule cell layer (gcl) (**c**), misplaced Purkinje cells (green) and abnormal Purkinje cell clusters (**d**); ml, molecular layer; scale bars: 100 μm. Arrows indicate misplaced *rln*^*A/A*^ Purkinje cells. See also Supplementary Figure 2: toluidine blue staining indicates not only misplaced Purkinje cells, but also dispersed granule cells in the cerebellum of *rln*^*A/A*^ mice. (**a**-**d**) Four animals per genotype were used for the analyses.

### 2.3. Reduced Reelin mRNA and protein levels in rln^A/A^ mice

Levels of mRNA and protein in *rln*^*A/A*^ and *rln*^*+/+*^ brains were measured by quantitative real-time PCR and immunoblotting with Reelin antibody G10. At embryonic day 14, Reelin protein levels were similar in *rln*^*A/A*^ and *rln*^*+/+*^ mice, whereas the levels decreased in *rln*^*A/A*^ mice relative to *rln*^*+/+*^ littermates at embryonic days 16 and 19, and at post-natal days 7, 14, 45 and >300 (Fig. 4a, b). Levels of Reelin mRNA in *rln*^*A/A*^ animals were similar to levels in *rln*^*+/+*^ litter-mates at embryonic day 14 and at postnatal day 5, while the levels were decreased in *rln*^*A/A*^ mice at postnatal days 7, 9, 14, 45 and >300 (Fig. 4b).

**Figure 4.**
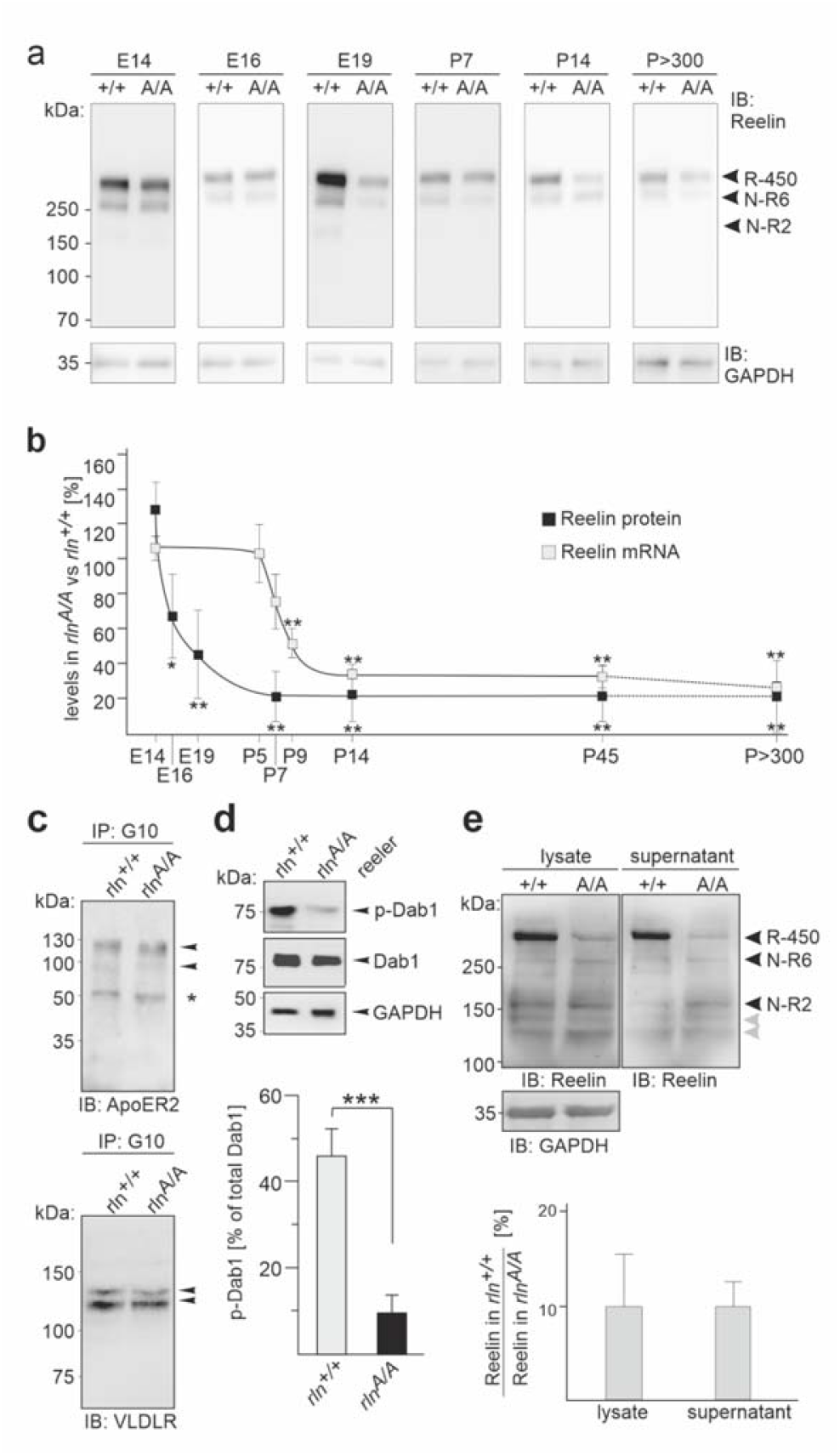
Reduced Reelin mRNA and protein levels and impaired Dab1 phosphorylation in *rln*^*A/A*^ mice. (**a**) Representative immunoblots of brain homogenates from wild-type (+/+) and mutant (A/A) mice with the Reelin antibody G10 and GAPDH antibody to control loading. (**b**) Quantification of full-length Reelin protein and Reelin mRNA levels in wild-type (*rln*^*+/+*^) and mutant (*rln*^*A/A*^) brains at different developmental stages. Mean values ± SD for Reelin protein and mRNA levels in *rln*^*A/A*^ brains relative to those in *rln*^*+/+*^ brains (six animals per genotype; *p<0.05, **p<0.01; one-way ANOVA with Bonferroni’s Multiple Comparison Test). (**c**) Co-immunoprecipitation of ApoER2 and VLDLR from brain homogenates of 16-day-old *rln*^*+/+*^ and *rln*^*A/A*^ embryos using Reelin antibody G10. Bands specific for ApoER2 and VLDLR are indicated by arrowheads; asterisk indicates non-specific bands. (**d**) Immunoblot analysis of brain homogenates from 16-day-old *rln*^*+/+*^ and *rln*^*A/A*^ embryos using an antibody against phosphorylated Dab1 (p-Dab1), total Dab1 and GAPDH to control loading. Levels of p-Dab1 were normalized to levels of total Dab1 protein. A repre-sentative immunoblot and mean values + SEM from 3 independent experiments are shown (***P<0.0001; ns, not significant; one-way ANOVA with Tukey’s Multiple Comparison Test). (**e**) Cell lysates were prepared and cell culture supernatants were collected from cortical neurons isolated from 16-day-old embryos and cultured overnight. They were subjected to protein precipitation and immunoprecipitation using Reelin antibody G10. The representative immunoblot shows secreted Reelin in the supernatants of cortical neurons from wild-type (+/+) and mutant (A/A) mice. Grey arrows indicate additional bands after an extended exposure of the immunoblots (these bands were not included for quantification). Quantification of Reelin levels in cell lysates and cell culture supernatants of cortical neurons from wild-type (+/+) and mutant (A/A) mice. Mean values + SD for the levels of full-length Reelin (R-450) in the lysates and supernatants (sup) of *rln*^*A/A*^ cultures relative to those in the lysates and supernatants (sup) of *rln*^*+/+*^ cultures (cultures from 3 animals per genotype; *p<0.001; one-way ANOVA with Bonferroni’s Multiple Comparison Test). Note: the low levels of secreted mutated Reelin correspond to the low levels of mutated Reelin in cell lysates.

To study whether mutated Reelin is associated with the canonical Reelin receptors ApoER2 and VLDLR, brain homogenates from *rln*^*A/A*^ and *rln*^*+/+*^ 16-day-old embryos were used for immunoprecipitation with Reelin antibody G10. Immunoprecipitates from *rln*^*A/A*^ and *rln*^*+/+*^ homogenates were positive for ApoER2 and VLDLR (Fig. 4c). When *rln*^*A/A*^ and *rln*^*+/+*^ homogenates were probed for phosphorylated Dab1, levels of phosphorylated Dab1 in *rln*^*A/A*^ were reduced (Fig. 4d). These results suggest that wild-type and mutated Reelin bind to ApoER2 and VLDLR, but that the stimulation of Dab-1 phosphorylation is impaired.

We further analyzed whether secretion of Reelin is impaired by the S/A_1283_ mutation. Supernatants of cultured cortical neurons from *rln*^*A/A*^ and *rln*^*+/+*^ mice were subjected to protein precipitation. Immunoblot analysis of protein precipitates and cell lysates showed that not only wild-type Reelin, but also S/A_1283_-mutated Reelin were detected in the cell supernatants and lysates (Fig. 4e). Moreover, the levels of S/A_1283_-mutated full-length Reelin were considerably lower relative to those of wild-type Reelin in cell supernatants and lysates (Fig. 4e). These results indicate that expression of mutated Reelin is low in *rln*^*A/A*^ mice, but that the mutation does not affect secretion of S/A_1283_-mutated Reelin.

### 2.4. Rescued migration of newborn rln^A/A^ cerebellar neurons by ectopic administration of wild-type Reelin, but not by S/A_1283_-mutated Reelin

The cerebellum of the *reeler* mouse shows the most obvious malformation when compared to other brain regions such as the hippocampus, cortex or olfactory bulb. Purkinje cells and cerebellar granule cells are dispersed and do not form compact layers. In our previous study [20] wild-type and *reeler* cerebellar explants showed increased migration of cerebellar granule cells out of the explant core after treatment with purified Reelin. Moreover, the cerebellar granule cells show positive *in situ* hybridization signals not only for Reelin’s binding partner cell adhesion molecule L1, which promotes neuronal migration, but also for Reelin and the components of the canonical Reelin signaling cascade Apo-ER2 (also known as LRP8), VLDLR, and Dab1 (Supplementary Figure 3a-e). Based on these observations, we asked whether purified S/A_1283_-mutated and wild-type Reelin would differentially affect migration of newly generated cerebellar granule cells. Since POMC-GFP has been found in newly generated immature granule cells [30] and has been used to study migration of individual neurons and their leading and trailing processes [31], we cross-bred *POMC-GFP* [30] and *rln*^*A/A*^ mice and used *rln*^*+/+*^*-POMC-GFP* and *rln*^*A/A*^*-POMC-GFP* offsprings to study migration of immature cerebellar granule cells. We observed that POMC-GFP is also expressed by newly generated cerebellar granule cells and have therefore prepared cerebellar explants from *rln*^*A/A*^*-POMC-GFP* neonatal cerebella for treatment with wild-type Reelin or S/A_1283_-mutated Reelin. Untreated *rln*^*A/A*^*-POMC-GFP* explants served as a control. The migration of the GFP-positive cerebellar granule neurons out of explants was monitored by fluorescence microscopy. Wild-type Reelin enhanced the speed of migration in comparison to untreated neurons (Fig. 5a, c, d; Supplementary Movie 2 and 4), whereas neurons treated with purified mutated Reelin did not increase the speed relative to untreated neurons (Fig. 5b, c, d; Supplementary Movie 2 and 3). These findings indicate that promotion of neuronal migration by Reelin depends on serine 1283.

**Figure 5.**
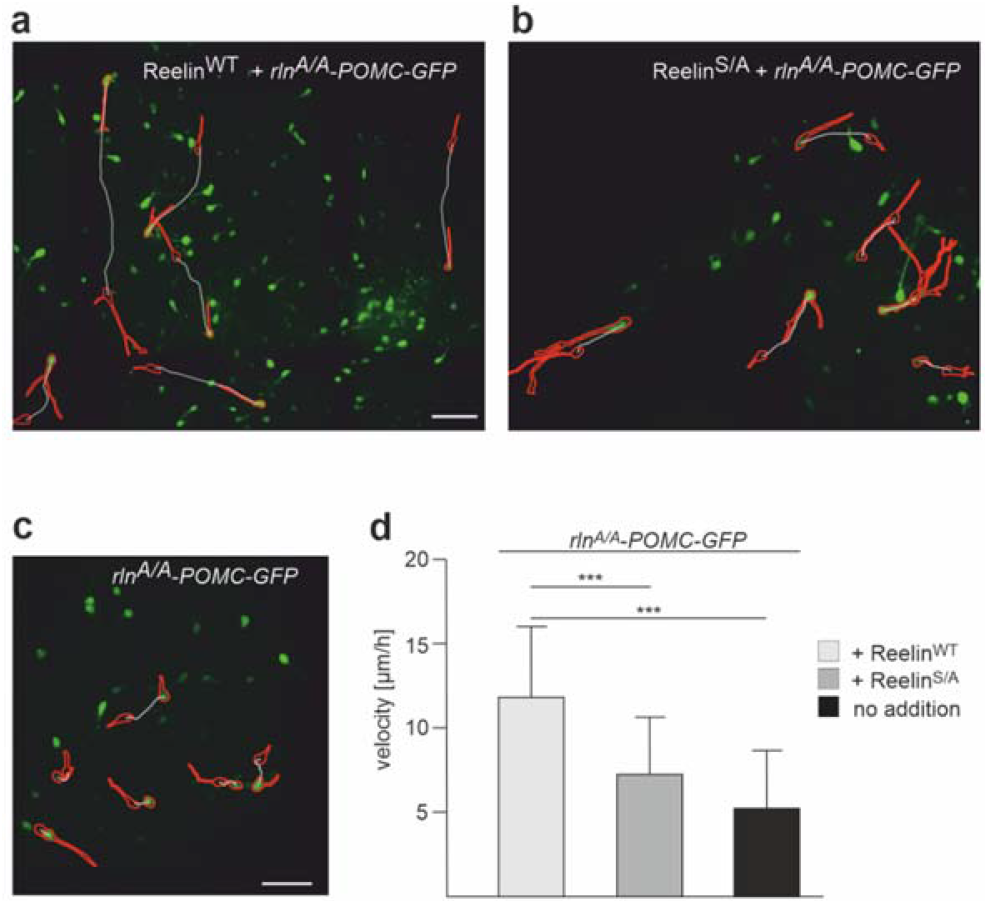
Real-time imaging of migrating newborn *rln*^*A/A*^ neurons after treatment with wild-type Reelin or S/A_1283_-mutated Reelin. Images of migrating newborn *rln*^*A/A*^-*POMC-GFP* cerebellar granule neurons within the explant area after treatment with heparin-purified wild-type Reelin (Reelin^WT^) (**a**) or S/A_1283_-mutated Reelin (Reelin^S/A^) (**b**) or without treatment (no addition) (**c**). (**d**) Quantification of the migration speed in μm/h of newborn POMC-GFP-positive neurons in *rln*^*A/A*^ explants treated with wild-type or mutated Reelin and with vehicle only. Mean values + SD from 4 independent experiments and at least 12 explants per treatment and experiment are shown (***p<0.0001; one-way ANOVA with Tukey’s Multiple Comparison Test, Brown-Forsythe Test and Bartlett’s Test). The migratory routes of six selected POMC-GFP granule cells are visualized. The shape of the selected cells at their initial and final positions is shown in red, and the migration trajectories of individual cells are indicated with white lines. See also Supplementary Movie 2-4 corresponding to the conditions from panels 4a, 4b, 4c.

### 2.5. Introduction of N-R6 into the rln^A/A^ cerebral cortex in utero normalizes neuronal migration

N- and/or C-terminal cleavage of Reelin by ADAMTS-4 yields fragments that contain either the N-terminal part only (N-R2), the N-terminal and central parts (N-R6), the central part only (R3-6), the central and C-terminal parts (R3-8), or the C-terminal part only (R7-8) [32] (for a schematic representation of the Reelin structure, see Fig. 7a). Since N-R6 was reported to be involved in proteolysis of L1 and L1-dependent cell migration [25], we investigated whether this fragment would rescue the aberrant migration of *rln*^*A/A*^ neurons. To this aim, a plasmid coding for N-R6 was introduced together with a CAG-GFP plasmid by *in utero* electroporation into the cerebral cortex of 13.5-day-old *rln*^*A/A*^ embryos. Sixty hours after electroporation, N-R6-expressing neurons were identified by GFP fluorescence, showing Reelin immunoreactivity. In comparison to the aberrantly migrating *rln*^*A/A*^ neurons electroporated with the CAG-GFP plasmid alone, a significant number of N-R6-expressing *rln*^*A/A*^ neurons had reached the upper cortical layers, as seen for *rln*^*+/+*^ neurons electroporated with CAG-GFP plasmid alone (Fig. 6a-c). The number of GFP-positive neurons per area was similar in the subventricular zone of *rln*^*+/+*^ mice and of *rln*^*A/A*^ mice electroporated with CAG-GFP plasmid encoding N-R6, while the number was decreased in *rln*^*A/A*^ mice electroporated with the empty plasmid in comparison to *rln*^*+/+*^ mice or *rln*^*A/A*^ mice electroporated with plasmid encoding N-R6 (Fig. 6d). These results suggest that N-R6 or its fragments promote the migration of cortical neurons.

**Figure 6.**
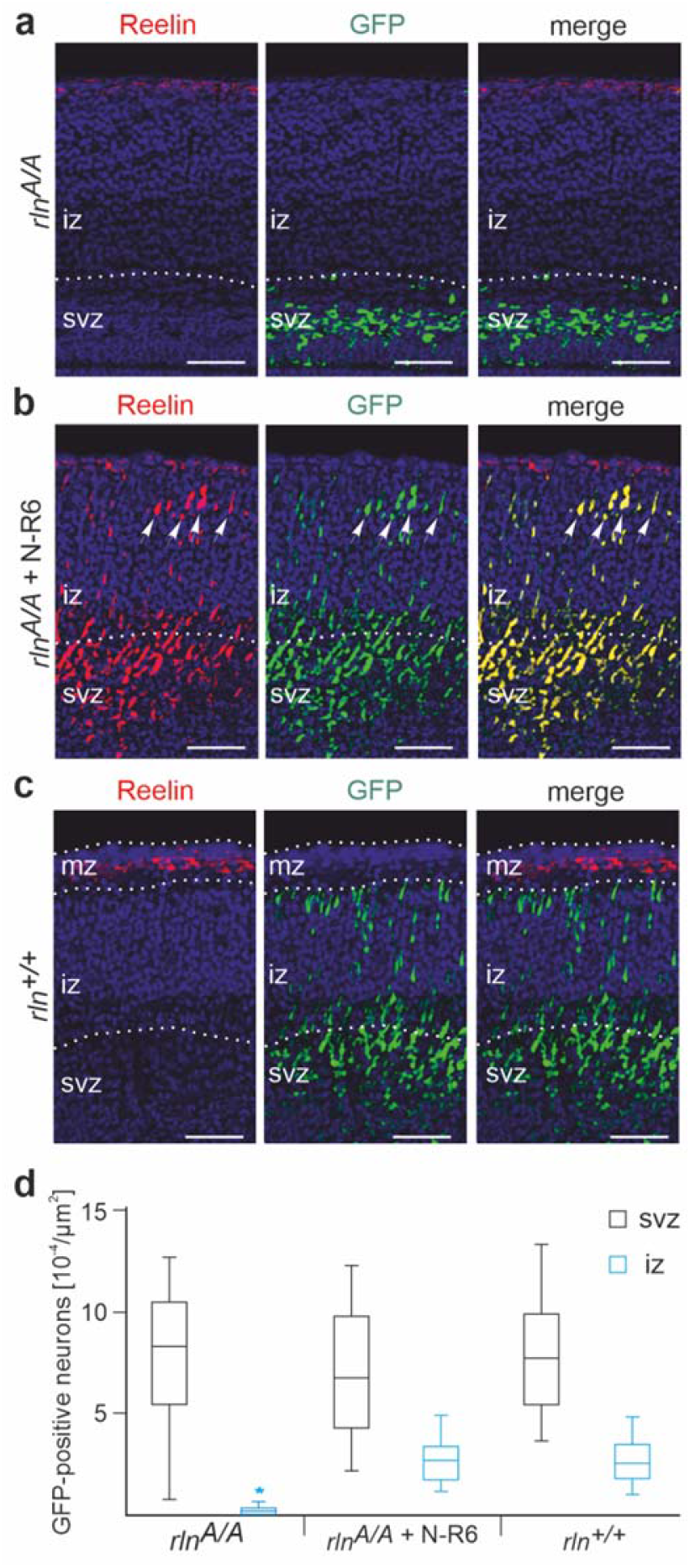
*In utero* electroporation of N-R6 into 13.5-day-old *rln*^*A/A*^ embryos promotes migration of cortical neurons and partially induces layer formation. (**a**) Sixty hours after *in utero* electroporation, many GFP-positive *rln*^*A/A*^ neurons (green) are retained in the subventricular zone (svz) of the *rln*^*A/A*^ cerebral cortex; white dashed line indicates the border between cortical plate and white matter. The border between svz and iz is identified by bright field light microscopy as a distinct border between callosal white matter and grey matter of the nascent cerebral cortex plate. (**b**) In contrast, in utero electroporation of N-R6 (*rln*^*A/A*^ + N-R6, electroporated neurons were identified by their immunoreactivity for G10 in combination with GFP expression) promotes neuronal migration from the svz towards the intermediate zone (iz) and towards a region close to the corresponding mz in the *rln*^*+/+*^ cerebral cortex. Note the formation of a rudimentary upper layer (white arrows) of *rln*^*A/A*^ neurons positive for both N-R6 (red) and GFP (green) near to the mz analogous region. (**c**) GFP-electroporated neurons from *rln*^*+/+*^ littermates were used for comparison. The *rln*^*+/+*^ mz contains many Reelin-expressing Cajal-Retzius cells (red) and the region underneath appears densely populated by electroporated neurons (green) at embryonic day 16.5. Nuclear staining with DAPI (blue); scale bars: 100 μm. (**d**) Brain sections from 4 electroporated mice per genotype (*rln*^*A/A*^: n=15 sections; *rln*^*A/A*^ + N-R6: n=18 sections; rln+/+: n=19 sections) were analyzed to determine the density of GFP-positive neurons in the subventricular zone (svz) (black) and intermediate zone (iz) (blue; * p<0.0001, two-way ANOVA). Note that few GFP-positive neurons are found in the iz of the *rln*^*A/A*^ cortex, indicating that *rln*^*A/A*^ neurons have migrated towards upper layers.

**Figure 7.**
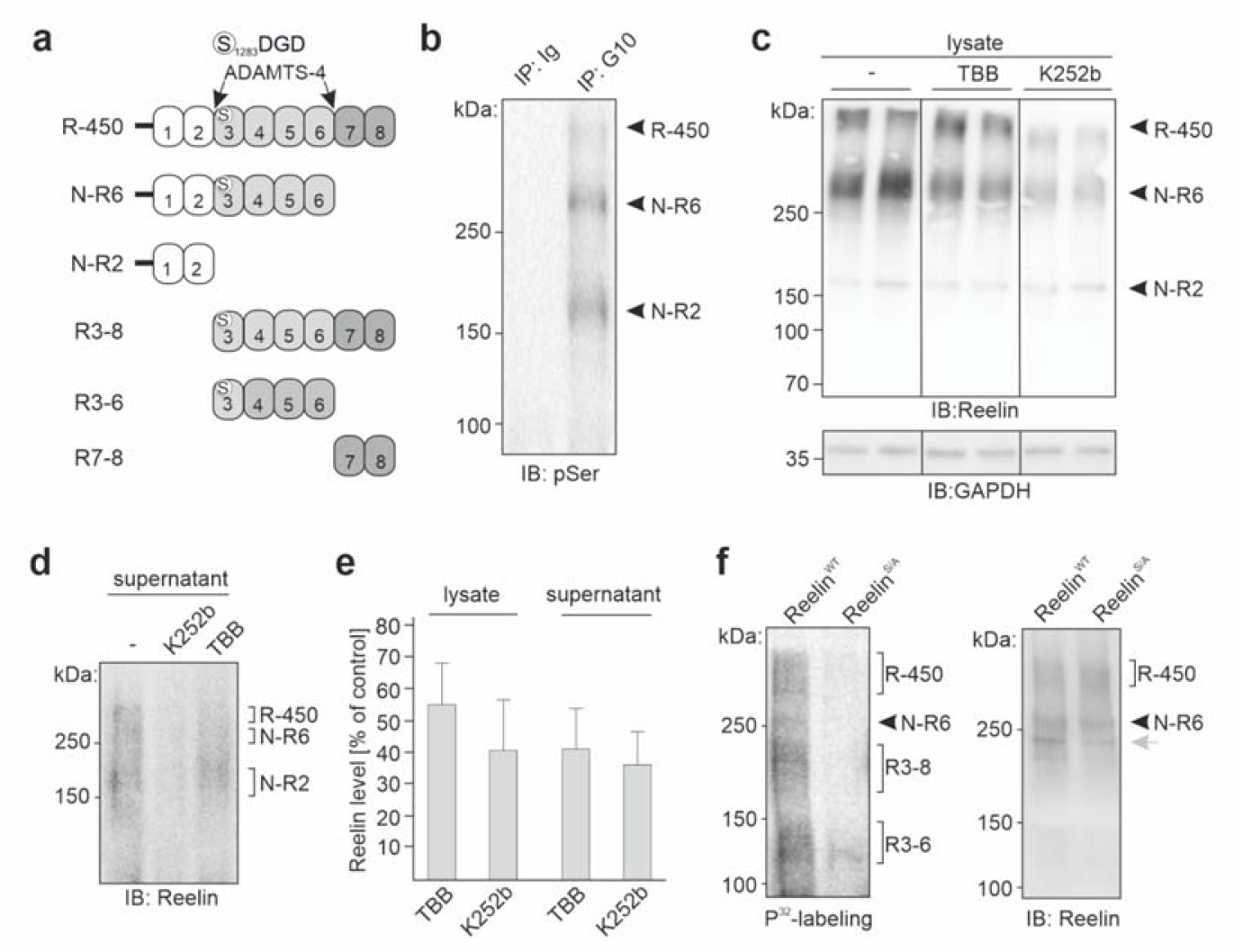
Reelin protein levels are reduced in cultured cerebellar neurons after inhibition of ecto-phosphorylation. (**a**) Schematic representation of the protein backbone of full-length Reelin (R-450) and its proteolytic fragments N-R6, N-R2, R3-8, R3-6 and R7-8 after N- and C-terminal processing by ADAMTS-4. Serine 1283 (circle) and the putative CK2 phosphorylation site SDGD in murine Reelin are indicated. (**b**) Supernatants of cultured cerebellar neurons were subjected to immunoprecipitation (IP) with Reelin antibody G10 or non-immune control antibody (Ig) followed by immunoblot analysis of the immunoprecipitates with a phospho-serine antibody (pSer). Full-length Reelin (R-450) and its fragments N-R6 and N-R2 are indicated. (**c, d**) Cultures of cerebellar neurons were treated with the CK2-specific inhibitor TBB or with the cell-impermeable broad-spectrum ecto-protein kinase inhibitor K252b. Cell lysates (**c**) and culture supernatants (**d**) were subjected to immunoblot analysis with Reelin antibody G10. GAPDH antibody was used to control loading of cell lysates. Representative immunoblots are shown. Lanes not adjacent to each other but from the same blot are separated by a vertical line. (**e**) Quantification of Reelin levels in cell lysates and cell culture supernatants. Mean values + SD from 4 independent experiments are shown for Reelin levels after TBB or K252b treatment relative to control treatment (set to 100%) (*p<0.05; Kruskal-Wallis One Way Analysis of Variance on Ranks with Dunn’s Multiple Comparison Test). (**f**) Wild-type Reelin (Reelin^WT^) and S/A_1283_-mutated Reelin (Reelin^S/A^) were subjected to in vitro phosphorylation using CK2 and radioactive ATP followed by blotting and analysis using a Phosphor-Imager. Diffuse signals and one distinct band with radiolabeled, phosphorylated proteins were observed (left panel). Full-length Reelin (R-450) and its serine 1283-containing fragments N-R6, R3-6 and R3-8 are indicated. Similar amounts of wild-type Reelin (Reelin^WT^) and S/A_1283_-mutated Reelin (Reelin^S/A^) were detected by immunoblot analysis with Reelin antibody G10 (right panel), showing that equal amounts of wild-type and mutated Reelin were used for in vitro phosphorylation. A grey arrow points to an additional band, which could not be assigned to one of the known Reelin forms.

### 2.6. Inhibition of ecto-phosphorylation of Reelin leads to reduced Reelin protein levels in cultured cerebellar neurons

The Reelin sequence S_1283_DGD represents a S/T-D/E-X-E/D motif for phosphorylation by CK2 [27]. This potential phosphorylation site for CK2 is present in the Reelin fragments N-R6, R3-6 and R3-8, but not in N-R2 and R7-8 (Fig. 7a). CK2 is not only active intracellularly, but also extracellularly. Extracellular CK2 (ecto-CK2) [33, 34] phosphorylates amyloid precursor protein and CD98 in their ectodomains [35, 36], C9 complement [37] and the extracellular matrix proteins vitronectin [38], collagen XVII [39], laminin [40], osteopontin and sialoprotein [41]. Ecto-CK2 is important for regulating ectodomain shedding, cell adhesion, migration and signaling as well as protein stability and degradation [42-44]. Based on these findings, we hypothesized that serine 1283 is phosphorylated by ecto-CK2 and that mutation of this phosphorylation site alters the structure and/or decreases the stability of Reelin, leading to enhanced internalization and degradation of Reelin.

To test this hypothesis, cerebellar neuron cultures, which contain mainly granule neurons (∼95%) that express Reelin [45-47] (Supplementary Figure 3b) and only low amounts of Reelin-expressing Purkinje cells (<0.1%) and interneurons (<5%), were treated with the CK2-specific inhibitor TBB [36, 39] and with K252b, which acts as a cell-impermeable broad-spectrum ecto-protein kinase inhibitor and inhibits ecto-CK2 [36, 39-41, 48, 49]. We first examined whether extracellular Reelin is phosphorylated by subjecting cell culture supernatants to immunoprecipitation with Reelin antibody G10 or non-immune control antibody and probing the immunoprecipitates with a phospho-serine-specific antibody. In the G10 immunoprecipitates, but not in the control immunoprecipitates, three major phospho-serine-positive bands were seen in the expected molecular weight range of the Reelin bands, thus indicating that extracellular full-length Reelin (R-450) and its fragments N-R6 and N-R2 are phosphorylated (Fig. 7b). We then treated cultured neurons with TBB and K252b and observed reduced levels of G10-positive Reelin bands in cell lysates and cell culture supernatants upon treatment when compared to non-treated neurons (Fig. 7c-e). Since intracellular Reelin is in secretory vesicles and cannot meet CK2 in the cytoplasm and since Reelin is internalized via its receptors, these results suggest that inhibition of ecto-phosphorylation of Reelin by CK2 results in either enhanced extracellular degradation, decreased internalization or increased internalization and intracellular degradation of non-phosphorylated Reelin. Of note, neither TBB nor K252b affected on the viability of neurons (TBB vs untreated: 100.8±4.9%; K252b vs untreated: 103.8±5.2%; mean values ± SEM from 3 independent experiments performed in duplicates; p> 0.05 in Kruskal-Wallis test with post-hoc Dunn’s multiple comparison test), indicating that the inhibitors are not toxic.

To support the notion that S_1283_DGD represents a site for CK2 phosphorylation and that serine 1283 is phosphorylated by CK2, we performed an *in vitro* phosphorylation assay using radioactive ATP, recombinant CK2 and purified wild-type and mutated Reelin. Three diffuse regions with phosphorylated bands and one distinct phosphorylated band were observed with wild-type Reelin, whereas only one faint band was seen with mutated Reelin (Fig. 7f). The immunoblot of purified wild-type and mutated Reelin with Reelin antibody G10 showed similar amounts of full-length Reelin and its G10-positive fragment N-R6 (Fig. 7f). According to the apparent molecular weights and since full-length Reelin (R-450) and its fragments N-R6, R3-6 and R3-8 contain serine 1283, we assume that the diffuse phosphorylated band regions correspond to differently glycosylated R-450, R3-8 and R3-6 and that the distinct band corresponds to N-R6. This result suggests that serine 1283 can be phosphorylated by CK2.

### 2.7. No alterations in the proteolytic processing of cell adhesion molecule L1 in brains of rln^A/A^ mice

Based on our finding that Reelin proteolytically cleaves L1 [25], we examined proteolytic L1 fragments in brain homogenates from *rln*^*+/+*^ and *rln*^*A/A*^ mice by immunoblot analysis using a L1 antibody against the intracellular domain. Levels of full-length L1 and of the 70-80 kDa L1 fragment that represents L1-70 generated by proteolysis via myelin basic protein (MBP) [49] were not altered in homogenates from *rln*^*A/A*^ mice in comparison to those in *rln*^*+/+*^ mice (Fig. 8a), indicating that the S/A_1283_ exchange does not suppress proteolytic cleavage of L1 *in vivo*.

**Figure 8.**
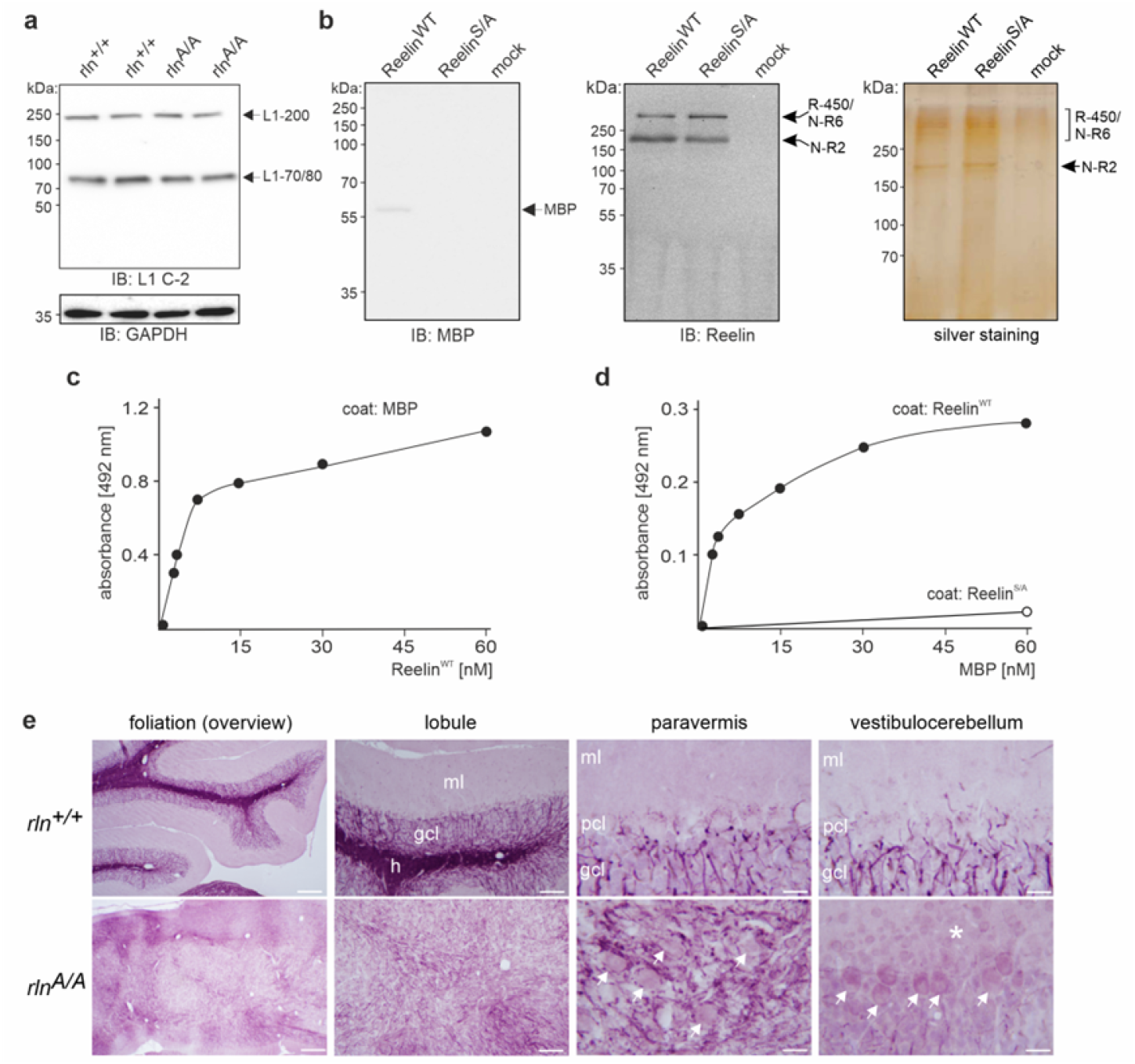
Proteolytic processing of L1 is not altered in mice expressing S/A_1283_-mutated Reelin but is mediated in vitro by the L1-cleaving protease MBP which binds to purified wild-type Reelin. (**a**) Immunoblot analysis of brain homogenates from 14-day-old *rln*^*+/+*^ and *rln*^*A/A*^ mice with the L1 antibody C-2 and GAPDH antibody to control loading shows unaltered proteolytic processing of L1 in *rln*^*A/A*^ brains in comparison to *rln*^*+/+*^ brains. Arrows indicate full-length L1 (L1-200) and the 70-80 kDa L1 fragment (L1-70/80). (**b**) Immunoblot analysis of heparin-purified wild-type Reelin (Reelin^WT^) or S/A_1283_-mutated Reelin (Reelin^S/A^) and of a mock control using pan-MBP antibody shows co-purification of MBP with wild-type Reelin, but not mutated Reelin (left panel). The arrow indicates the proteolytically active MBP form of ∼60 kDa. Immunoblot with Reelin antibody G10 (middle panel) and silver staining (right panel) show that similar amounts wild-type Reelin (Reelin^WT^) and S/A_1283_-mutated Reelin (Reelin^S/A^) were used. Full-length Reelin (R-450) and its fragments N-R6 and N-R2 are indicated. (**c, d**) ELISA using purified MBP or heparin-purified wild-type Reelin (Reelin^WT^) or S/A_1283_-mutated Reelin (Reelin^S/A^) as substrate coat or soluble ligand shows concentration-dependent binding of wild-type Reelin to substrate-coated MBP (**c**) and concentration-dependent binding of MBP to substrate-coated wild-type Reelin, but not to substrate-coated mutated Reelin (**d**). (**e**) Horizontal cerebellar sections from one-month-old *rln*^*+/+*^ and *rln*^*A/A*^ mice immunostained with an MBP antibody show MBP-positive *rln*^*+/+*^ white matter with a densely stained hilar region and heavily myelinated fibers within a compact granule cell layer with traversing MBP-positive myelinated fibers reaching the Purkinje cell monolayer and wrapping around Purkinje cell somata, and a prominent and myelin-free molecular layer. In contrast, the *rln*^*A/A*^ cerebellum displays a severely distorted MBP-stained white matter with dispersed myelin fibers and no layers. The MBP immunostaining signal in the *rln*^*A/A*^ cerebellum appears to be weaker than in the *rln*^*+/+*^ cerebellum. Overview, scale bars: 300 μm; lobule, scale bars: 100 μm; paravermal and paramedical lobule, scale bars: 30 μm. Arrows indicate dispersed *rln*^*A/A*^ Purkinje cells, also at the pia; * indicates the pial surface where cerebellar granule cells are found; gcl: granule cell layer, h: hilar region, ml: molecular layer, pcl: Purkinje cell layer. Four animals per genotype were used for analysis.

### 2.8. Binding of Reelin to MBP via serine 1283

We have shown that treatment of immunopurified L1 with heparin-purified wild-type but not S/A_1283_-mutated Reelin generates of a 70-80 kDa L1 fragment [25]. However, here we showed that proteolytic cleavage of L1 is not altered in *rln*^*A/A*^ mice, which display very low levels of Reelin, suggesting that L1 is not cleaved by Reelin. Thus, we assumed that in our previous *in vitro* experiments the heparin-purified wild-type Reelin but not the mutated Reelin may have contained a co-purifying protease which cleaves L1. We had identified a proteolytically active MBP form (60 kDa sumoylated dynamin-MBP chimera) as a serine protease which cleaves L1 [50], thereby generating the fragment L1-70 [25, 51]. We thus tested by immunoblot analysis whether MBP co-purifies with heparin-purified wild-type Reelin. Indeed, MBP was detected as a ∼60 kDa band by the pan-MBP antibody in the sample with wild-type, but not mutated Reelin (Fig. 8b). This result indicates that the proteolytically active ∼60 kDa MBP form co-purifies with wild-type, but not mutated Reelin. The findings also suggest that MBP co-purifies with wild-type Reelin and is responsible for the generation of the 70-80 kDa fragment in the previous *in vitro* studies.

The co-purification of MBP with wild-type, but not mutated Reelin suggests that MBP binds to wild-type, but not mutated Reelin. To substantiate this notion, we performed ELISA with MBP and either purified wild-type or mutated Reelin. Wild-type Reelin bound to substrate-coated MBP in a concentration-dependent and saturable manner (Fig. 8c). A similar concentration-dependent and saturable binding of MBP to substrate-coated wild-type Reelin was observed, whereas MBP did not bind to substrate-coated mutated Reelin even at high concentrations (Fig. 8d). These results indicate that MBP binds to wild-type Reelin and that this binding depends on serine 1283.

Based on these findings, we asked whether Reelin would affect the expression or/and tissue localization of MBP. Immunostaining for MBP in *rln*^*+/+*^ and *rln*^*A/A*^ cerebellar slices was performed, since MBP is expressed in cerebellar neurons, including granule cells (Supplementary Figure 3f). In *rln*^*+/+*^ cerebella MBP was characteristically present in white matter with densely stained hilar region containing well-myelinated fibers, compact granule cell layer with traversing myelinated fibers reaching the Purkinje cell layer and ensheathing Purkinje cell somata, and a prominent and myelin-free molecular layer (Fig. 8e). In contrast, in *rln*^*A/A*^ cerebella the MBP-positive white matter is severely distorted with dispersed myelin fibers and lack of layers (Fig. 8e). Of note, the MBP staining intensity in the *rln*^*A/A*^ cerebellum was weaker than in the *rln*^*+/+*^ cerebellum (Fig. 8e).

## 3. Discussion

The lack of Reelin expression in *reeler* mice has been shown to cause abnormal neuronal migration, impaired motor coordination, tremors, and ataxia [3], thus indicating that Reelin is required for proper brain development and functioning. Here, we provide evidence that mice expressing Reelin with mutated serine 1283 show a *reeler*-like phenotype with abnormal neuronal migration, disrupted layering in the cerebral cortex, hippocampus and cerebellum, and strong motor deficits accompanied by tremor and ataxia as well as spontaneous epileptic seizures which are not shown in *reeler* mice. *In vitro* experiments showed that S/A_1283_-mutated Reelin does not promote neuronal migration in contrast to wild-type Reelin, indicating that serine 1283 in Reelin is required for Reelin-dependent neuronal migration.

Previous *in vitro* experiments by us and others using cells expressing Reelin with mutated serine 1283 (S/A or S/C) [25, 52] showed that these mutated proteins are secreted, indicating that mice carrying the S/A_1283_ mutation are not abnormal because of intracellular retention of Reelin. Of note, in the *reeler* mouse carrying an Orleans mutation lacking 73 amino acids of the C-terminus, Reelin cannot be secreted by the cells [53, 54]. Here, we confirm that the S/A_1283_ mutation does not affect Reelin’
ss secretion. Moreover, the presence of the N-R6 fragment in *rln*^*A/A*^ brains indicates that the S/A_1283_-mutated full-length Reelin is released into the extracellular space where it is cleaved at its C-terminus between Reelin repeat 6 and 7 by extracellular serine proteases and metalloproteinases [20, 55-57].

The serine 1283-carrying N-R6 fragment contains the N-terminal region which is involved in Reelin oligomerization and has regulatory functions in canonical Reelin signaling [19, 32, 58-62]. Importantly, *in utero* electroporation of *rln*^*A/A*^ mice with N-R6 rescued the impaired migration of neurons. Reelin has been shown to alter neuronal migration when expression is engineered to occur in regions where Reelin is normally not present [63, 64]. Indeed, N-R6 expression in subventricular neurons promotes their migration towards the marginal zone. It is therefore likely that the serine 1283-containing N-R6 fragment is crucial for neuronal migration and thus for brain development and functioning.

We would like to point out that it is difficult to dissect the functions of N-R6 from the functions of full-length Reelin in the wild-type tissue. However, it seems that expression of N-R6 affects neuronal migration: impaired processing of Reelin and accumulation of unprocessed Reelin are associated with granule cell dispersion in experimental models of epilepsy [55]. Interestingly, the proteolytic generation of N-R6 was inhibited by TIMP1, an inhibitor of matrix metalloproteinase activity. This inhibitor was increased in hippocampal slices with kainate-induced epileptiform activity [55]. Thus, the epileptic seizures seen in *rln*^*A/A*^ mice might be the consequence of increased TIMP1 levels and reduced N-R6 levels.

It is presently not known, whether Reelin fragments have different or even opposing functions. For instance, the central Reelin fragment activates signaling via the cognate Reelin receptors. We here show that N-R6 promotes migration. One could hypothesize that the central Reelin fragments may act as stop signals, whereas the fragments that contain the N-terminus, e.g. N-R6 and its fragments, may promote migration. Moreover, Reelin overexpression by *in utero* electroporation seems to stop, rather than promote, migration [65, 66]. We hypothesize that overexpression of Reelin could lead to accumulation of unprocessed Reelin or processed central Reelin fragments which would then act as a stop signal as seen by granule cell dispersion in the experimental models of epilepsy [55].

Although *in utero* electroporation introduces Reelin into cells where it is normally not expressed at early developmental stages, we used this system to test whether Reelin may help Reelin-deficient neurons to migrate normally. Interestingly, electroporation of N-R6 into *rln*^*A/A*^ embryos produced N-R6-expressing neurons in the intermediate zone of the cerebral cortex, indicating that some neurons have migrated further than others. Moreover, only very few GFP-expressing *rln*^*A/A*^ neurons were found in the upper cortical layers of *rln*^*A/A*^ embryos which were electroporated with GFP alone. This result suggests that at later developmental stages more *rln*^*A/A*^ neurons migrate to upper layers, as seen for *reeler* neurons 5 to 7 days after *in utero* electroporation [67, 68]. Unfortunately, in multiple attempts we failed to electroporate full-length Reelin and full-length S/A_1283_-mutant Reelin *in utero*. A reason for this could be that full-length Reelin is very large (10383 nucleotides). However, even if it were electroporated successfully, full-length Reelin might be proteolytically cleaved, thus preventing to dissociate the effects of full-length Reelin from its fragments. Furthermore, the N-terminal cleavage site in Reelin is so close to serine 1283 that fragments N-R2, R3-6 and R3-8 resulting from this cleavage may affect S_1283_, a hypothesis that would be worthwhile to follow up.

Immunoprecipitation suggests that S/A_1283_-mutated Reelin binds to ApoER2 and VLDLR, whereas it only marginally triggers Dab1 phosphorylation in mutant mice. Thus, we propose that the mere binding of mutated Reelin to ApoER2 and VLDLR is not sufficient to induce proper phosphorylation of Dab1 and activation of canonical Reelin signaling.

Since serine 1283 is a part of a potential CK2 phosphorylation site and since Reelin is phosphorylated by ecto-CK2, it is conceivable that ecto-phosphorylation contributes to Reelin’s role in neural development and function. In mice expressing Reelin with disrupted CK2 phosphorylation site at serine 1283, Reelin protein expression decreases at approximately embryonic day 16, reaches lowest values already at postnatal day 7 and continues to stay low at later stages. We propose that impaired ecto-phosphorylation of serine 1283 affects the structure and/or stability of Reelin, probably leading to enhanced degradation and internalization via cognate cell surface receptors. It is conceivable that impaired signaling due to defects in CK2-mediated ecto-phosphorylation of Reelin leads to migratory deficits [69], such as ‘hypermigration’ of late-generated neurons. The levels of brain Reelin mRNA decrease around postnatal day 7, reach the lowest values around postnatal day 14 and continue to remain low thereafter. The delayed decrease of mRNA levels relative to protein levels indicates that reduced protein levels lead to a decrease in mRNA levels, possibly owing to a feedback mechanism. With this hypothesis, we assume that reduced levels of Reelin may contribute to the phenotype of *rln*^*A/A*^ mice. Nonetheless, we cannot exclude that kinase substrates other than Reelin might be involved in regulating the changes in Reelin levels.

Since the generation of transgenic mice via zinc finger nucleases can lead to site-off effects, we confirmed by different methods that the mutation was targeted specifically to the *Reelin* gene. We were therefore pleased that the S/A_1283_ point mutation led to a *reeler*-like phenotype that resembles but does not copy that of *reeler* mice. The pronounced loss of body size and weight in *rln*^*A/A*^ mice is usually not observed in *reeler* mice. In addition, *reeler* mice do not show spontaneous epileptic seizures [70] as seen in *rln*^*A/A*^ mice (Supplementary Movie 1) and show increased numbers of Ca-jal-Retzius cells [71] in contrast to decreased numbers in *rln*^*A/A*^ mice. These differences suggest that *rln*^*A/A*^ mice are not simply functional knockout mice.

Since mutant Reelin is produced at RNA and protein level and then rapidly degraded at protein level, we did not investigate possible errors in the mechanisms of Reelin’s synthesis, neither at translational nor transcriptional levels. Yet, we cannot exclude such phenotypes, thus calling for mutations produced by different genome editing techniques. It must also be considered that the specific amino acid substitution serine 1283 may determine the mutant phenotype. Replacement of serine 1283 by cysteine [52] could hinder proteases to degrade Reelin, since cysteine contains a thiol group that can form disulfide bridges inducing conformational changes in protein structure. It also cannot be excluded that this replacement leads to structural stabilization in contrast to the S/A_1283_ exchange, where alanine might lead to destabilization of Reelin.

It needs to be emphasized that the degradation of mutated Reelin in the present mouse model occurs rapidly between embryonic day 14 and 19, leading to 50% less protein and low protein levels can be detected from postnatal day 7 onwards. Hence, this degradation could contribute to the *reeler*-like phenotype of the *rln*^*A/A*^ mice.

It is noteworthy in this respect that full-length S/A1_283_ mutated Reelin is not degraded HEK293 cells. In tissue explants, mutated and inactive Reelin therefore may not stimulate migration, possibly due to proteases in the explants that are rapidly degrading it.

It has been reported that Reelin is a protease and functions as a serine protease and that serine 1283 is the active site for Reelin-mediated proteolysis [24-26]. Another study could not show Reelin’s proteolytic activity biochemically [52]. This led to the debate whether Reelin is a serine protease and, more importantly even, whether this proteolytic activity is important for brain development. Here, we provide evidence that Reelin does not cleave its interaction partner L1 *in vivo*, thus contradicting the notion that Reelin is a protease. However, one could hypothesize that Reelin associates with serine proteases, such as L1-cleaving MBP, and that Reelin’
ss serine 1283 and its phosphorylation are important for these interactions and for regulating the activity of associated proteases. These putative Reelin-associated serine proteases could be responsible for the cleavage of certain proteins in *in vitro* biochemical proteolysis assays. One could ask whether MBP alone can act as Reelin’s protease. MBP is much later expressed than Reelin, so that both molecules would only join at later ontogenetic stages, after the end of neuronal migration, indicating that the interaction between MBP and Reelin is not involved in cerebral cortex development. As to possible roles in late developing structures such as the cerebellum as well as the fiber tract systems and their myelination, this unconventional interaction calls for further experimental evidence.

In summary, our study elucidates the potential roles of Reelin fragment N-R6 in the navigation of migrating neurons *in vivo* and suggests that serine 1283 is important for Reelin’s functions in signaling and brain development, but not proteolysis.

## 4. Materials and Methods

For detailed descriptions of methods see Supplementary Methods.

### 4.1. Animals

The *CXCR4-GFP* mouse [28, 29] and *POMC-GFP* mouse [30] have been described. Animals were housed at 22°C on a 12 h light/12 h dark cycle with ad libitum access to food and water. For staging of embryos, the day of the vaginal plug was considered embryonic day 0.5. The *rln*^*A/A*^*-POMC-GFP* and *rln*^*A/A*^*-CXCR4-GFP* mice were generated by cross-breeding of *rln*^*A/A*^ mice with *POMC-GFP* or *CXCR4-GFP* mice. All experiments were conducted in accordance with the German and European Community laws on protection of experimental animals, and all procedures used were approved by the responsible authorities of the State of Hamburg (Behörde für Wissenschaft und Gesundheit, Amt für Gesundheit und Verbraucherschutz, Lebensmittelsicherheit und Veterinärmedizin; animal permit numbers ORG-679, ORG-604, ORG-578, TVA 6/14, TVA 45/14) and the State of Nordrhein-Westfalen (Landesamt für Natur, Umwelt und Verbraucherschutz Nordrhein-Westfalen; animal permit numbers 81-02.04.2018.A306 and 81-02.04.2019.A308). The manuscript was prepared following the ARRIVE guidelines for animal research.

### 4.2. Antibodies and reagents

Mouse monoclonal Reelin antibody G10 (Millipore, MAB5364) directed against N-terminal epitopes and rabbit p-Ser491 Dab1 antibody (AB9642) were from Merck Chemicals (Darmstadt, Germany). Rabbit polyclonal Brn2 antibody (sc-28594), mouse monoclonal L1 antibody C-2 (NCAM-L1; sc-514360), goat polyclonal antibody against MBP (sc-13914) and mouse monoclonal antibody against phosphorylated serine (phospho-Ser) (sc-81514) were from Santa Cruz Biotechnology (Heidelberg, Germany). Rabbit polyclonal calbindin antibody (C8666) and mouse monoclonal NeuN antibody (clone A60; Millipore, MAB377B) were from Sigma-Aldrich (Taufkirchen, Germany). Rabbit polyclonal Cux1 antibody (homeobox protein cut-like 1 antibody; ab140042) rabbit polyclonal ApoER2 antibody (ab52905,) and rabbit polyclonal MBP antibody (ab40390) were from Abcam (Cambridge, UK). Rabbit GAPDH antibody (#2118) was from Cell Signaling Technology Europe (Frankfurt am Main, Germany). Goat polyclonal VLDLR antibody (AF2258) was from R&D Systems (Minneapolis, MN, USA). Rabbit Dab1 antibody (100-401-225) was from Rockland (Limerick, PA, USA). Secondary donkey antibodies conjugated with Cy3, Cy2 or horseradish peroxidase were from Dianova (Hamburg, Germany). MBP purified from bovine brain (#6420-0100) was from BioRad (AbD Serotec; Puchheim, Germany). Primers were from Metabion International (Planegg, Germany). The CK2 inhibitor 4,5,6,7-tetrabromobenzotriazole (TBB; CAS 17374-26-4) was from Santa Cruz Biotechnology and K252b (CAS No: 99570-78-2) was from Biozol (Eching, Germany). Matrigel (Corning^®^ Matrigel^®^ Basement Membrane Matrix, LDEV-Free) was from Corning Life Sciences (Amsterdam, The Netherlands).

### 4.3. Generation of gene-edited mice, Southern blot analysis, genomic sequencing and genotyping

The wild-type Reelin sequence (NC_000071.6; Gene ID: 19699) was used for the design and screening of zinc finger nuclease (ZFN) pairs (Sigma-Aldrich). Linearized plasmids carrying cDNA encoding ZNF1 or ZNF2 were subjected to reverse transcription as described [51]. The ZFN transcripts and the plasmid carrying the 1,600 bp donor sequence and the mutation (Genewiz, Leipzig, Germany) were injected into the pronuclei of one-cell-stage embryos. Injection and implantation of pronuclei into foster mothers was performed as described [51]. In the mutated sequence, the TCA codon coding for serine 1283 was replaced by GCG (coding for alanine) generating thereby a restriction site for AciI.

The correct insertion of the mutation into the genomic DNA was confirmed by Southern blot analysis, whole genome sequencing and genomic sequencing.

For genotyping, the Phire Animal Tissue Direct PCR Kit (ThermoFisher Scientific), mouse tail biopsies, the primers P1 and P2, and AciI (New England Biolabs, Frankfurt am Main, Germany) were used.

### 4.4. Pencil grab test and assessment of general locomotion

One-month-old female *rln*^*+/+*^ and *rln*^*A/A*^ mice were subjected to the pencil grab test and were placed in empty cages or on the barred cage lid to assess stance, posture and coordination.

### 4.5. Isolation of RNA, reverse transcription and quantitative real-time PCR

For quantitative real-time PCR reverse transcribed brain RNA and Reelin-specific primers (5’- AAC TTC TAT GAG AAG CCA GCT TTC -3’; 5’- AAC TGC AGC ACA TAT CCA GGT TTC -3’) were used.

### 4.6. Plasmids and site-directed mutagenesis

The pcDNA3 plasmids with Reelin cDNA coding for full-length Reelin comprising amino acids 1-3461 [72] or coding for the Reelin fragment N-R6 comprising amino acids 1-271227 were kindly provided by T. Curran and A. Goffinet. For mutagenesis, a sequence region of 5,000 bp between SnaBI and AgeI restriction sites was amplified using the HiFi amplification kit (Clontech, Takara Bio Europe, Saint-Germain-en-Laye, France), the plasmid coding for full-length Reelin and the primers 5’-AAC CCC ACC TAC TAC GTA CCG GGA CAG GAA TAC-3’ and 5’-GT TAC TGC AAA CCG GTC TCC ATC CGC CTT TCC AAA CAC CAT GGC TG-3’. Using the In Fusion Cloning Kit (Clontech) the 5,000 bp amplicon carrying the mutation was inserted into the full-length Reelin-coding plasmid digested with SnaBI and AgeI.

### 4.7. In utero electroporation

*In utero* electroporation of *rln*^*A/A*^ E13.5 embryos using N-R6-coding plasmid (2 μg) and *CAG-IRES-GFP* vector (2 μg) pre-mixed with 0.01% Fast Green was performed as described [25]. GFP-expressing brains from electroporated *rln*^*A/A*^ embryos identified by genotyping were collected under a fluorescence microscope and fixed overnight followed by an overnight incubation in a sucrose solution.

For standard fluorescence microscopy, electroporated brains were embedded in agarose and sectioned on a vibratome. The 80 μm-thick slices were immersed in blocking solution, washed, and incubated overnight with Reelin antibody G10 (1:500). The tissue sections were washed, incubated overnight with secondary antibodies (1:250), counterstained with DAPI, and mounted in Mounting Medium (Dako, Agilent, Santa Clara, CA, USA).

### 4.8. Tissue preparation, immunohistology and imaging

Brain slices of 25 μm thickness were used for antigen demasking, blocking, antibody staining and imaging. After paraffin embedding of fixed brains sections of 10 μm were prepared, deparaffinized and subjected to immunohistochemistry with the following primary antibodies: Reelin antibody G10 (1:500), rabbit MBP antibody (1:500), NeuN antibody (1:1,000), and antibodies directed against Brn2 (1:250), calbindin (1:1,000) or Cux1 (1:250). Secondary donkey antibodies conjugated with Cy3 and Cy2 (1:500) and Fluoromount containing DAPI (Sigma-Aldrich) were used. Images were taken on a confocal fluorescence microscope and processed with ImageJ.

For toluidine blue staining and microscopical analysis, osmicated and fixed cerebella were embedded in Araldite resin and cut on an ultramicrotome. 0.75 μm-thick sections were collected and stained with toluidine blue.

For Nissl staining using cresyl violet and imaging, 10 μm-thick sagittal sections of paraffin-embedded brains were used.

### 4.9. Counting of Cajal-Retzius cells in mouse cerebral cortices

Mounted sagittal brain slices were subjected to fluorescence microscopy with focus on the marginal zone of the somatosensory cortex in *rln*^*+/+*^*-CXCR4-GFP* mice and on the corresponding analogous region in *rln*^*A/A*^*-CXCR4-GFP* mice. The marginal zone in the wild-type and its analog in the mutant mice was divided in four to five segments, each segment with a depth of 70 μm (from the pial surface) and width of 50 μm. Each segment was divided into seven bins of 10 μm depth. The Cajal-Retzius cells were counted in each bin and the CR cell distribution across the seven bins of each segment was assessed.

### 4.10. Preparation of brain homogenates, SDS-gel electrophoresis, silver staining, immunoblot analysis and immunoprecipitation

Preparation of brain homogenates, SDS-gel electrophoresis and immunoblot analysis using Reelin antibody G10 (1:1,000), L1 antibody C-2 (1:1,000), goat MBP antibody (1:1,000), phospho-Ser antibody (1:1,000), GAPDH antibody (1:2,000) and secondary antibodies coupled to horseradish peroxidase (1:10,000) was carried out as described [46].

For immunoprecipitation, cell culture supernatants of cultured cerebellar neurons were incubated overnight with Reelin antibody G10 and Protein A/G beads (Santa Cruz Biotechnologies) or brain homogenates were resuspended in immunoprecipitation buffer (25mM Tris-HCl, 150 mM NaCl, 1% Nonidet P-40, 1% sodium deoxycholate, 0.1% SDS, pH 7.6) and subjected to preclearing and overnight incubation with Reelin antibody G10 and Dynabeads Protein G (ThermoFisher Scientific). For silver staining of SDS-gels PierceTM Silver Stain for mass spectrometry (ThermoFisher Scientific) was used.

### 4.11. Purification of wild-type and S/A_1283_-mutated Reelin

Isolation of wild-type and mutated Reelin from transfected HEK cell culture supernatants and assessment of protein yield and purity were carried out as described [25, 52]. Supernatants from mock-transfected HEK cells served as controls.

### 4.12. Treatment of dissociated cell cultures and explants and imaging of migrating cerebellar granule cells

For treatment with CK2 inhibitors, cerebellar neurons were maintained in serum-free medium overnight. Neurons were then incubated for 2 h in fresh serum-free medium containing 25 μM TBB and 0.025 % DMSO, 500 nM K252b and 0.025 % DMSO or only 0.025 % DMSO. Cell culture supernatants were collected and centrifuged at 10,000 g and 4°C. Cells were lysed using RIPA buffer. Cell survival was determined using neutral red uptake assay after treatment of cerebellar neurons with 25 μM TBB and 500 nM K252b for 30 h.

Cell culture supernatants from cerebral cortex neurons were centrifuged at 10,000 g and 4°C and subjected either to protein precipitation or to immunoprecipitation using Reelin antibody G10 and Dynabeads Protein G (ThermoFisher Scientific).

Cerebellar explants were incubated with purified wild-type or S/A_1283_-mutated Reelin (5 ng), seeded onto Matrigel-coated coverslips, incubated for 3 h, and subjected to real-time microscopy and quantitative analyses.

### 4.13. In vitro CK2 phosphorylation assay using purified wild-type and mutated Reelin

Wild-type Reelin or S/A_1283_-mutated Reelin (1 μg) were incubated for 1 h at 30°C in 100 μl containing 100 μM ATP, 10 μCi ɣ-P32-ATP (Hartmann Analytics, Braunschweig, Germany), 500 U CK2 (New England Biolabs), and 1x NEB-uffer for Protein Kinases (New England Biolabs). The samples were then subjected to protein precipitation, SDS-gel electrophoresis and blotting onto nitrocellulose membrane. Radiolabeled proteins were visualized by a Phos-phor-Imager (Molecular Dynamics, Sunnyvale, USA).

### 4.14. ELISA

ELISA with substrate-coated MBP (250 ng) or purified wild-type or mutated Reelin (2.5 μg) and different concentrations of purified wild-type Reelin or MBP was performed as described20. Reelin antibody G10 (1:100), goat MBP antibody (1:100) and anti-mouse and anti-goat HRP-coupled secondary antibody (1:2,000) were used for detection of Reelin and MBP.

## Acknowledgments

The authors are grateful to D. Lutz, Dagmar Drexler, Bianka Brunne, E. Förster and M von Düring for their contributions to this study. We thank E. Kronberg, S. Deutsch, U. Wolters, L. Augustinowski, K. Rumpf, C. Wojczak, J. Willms and S. Homann for excellent technical support, A. Failla for the possibility to use the IMARIS software, M. Richter and F. Calderon de Anda for providing the CAG-IRES-GFP vectors, M. Hohlbaum and S. Hoffmeister-Ullerich for sequencing, and G. Korkmaz and J. Wang for whole genome sequencing. We are grateful to T. Curran and A. Goffinet for Reelin antibodies and plasmids, T. Pohlkamp for helpful comments, and G. Rune for the opportunity to use the Keyence microscope. We also thank G. Westbrook for providing the *POMC-GFP* transgenic mice. We are very grateful to Michael Frotscher for his deep interest in the Reelin project and invaluable support over many years. Michael Frotscher was supported by the Deutsche Forschungsgemeinschaft (FR 620/12-2 and FR 620/13-1).

## Data Availability

All data obtained and analyzed in the current study are available from the corresponding authors upon reasonable request.

## Author Contributions

Conceptualization, Gabriele Loers, Ralf Kleene and Melitta Schachner; Investigation, Gabriele Loers, Ralf Kleene, Ahmed Sharaf, Shaobo Wang, Hardeep Kataria, and Irm Hermans-Borgmeyer; Methodology, Max Anstötz, Irm Hermans-Borgmeyer; Resources, Max Anstötz; Visualization, Ralf Kleene; Writing – original draft, Gabriele Loers, Ralf Kleene; Writing – review & editing, Gabriele Loers, Ralf Kleene and Melitta Schachner.

## Abbreviations

ApoER2: apolipoprotein E receptor 2
CK2: casein kinase 2
Dab1: disabled homolog 1
GFP: green fluorescent protein
N-R2: Reelin fragment comprising the N-terminal domain and the Reelin repeats 1 and 2
N-R6: Reelin fragment comprising the N-terminal domain and the Reelin repeats 1 to 6
R3-6: Reelin fragment comprising the Reelin repeats 3 to 6
R3-8: Reelin fragment comprising the Reelin repeats 3 to 8
R7-8: Reelin fragment comprising the Reelin repeats 7 to 8
rln^A/A^: Reelin mutant mice carrying the mutation of serine 1283 to alanine
rln^+/+^: Wild-type littermates of the Reelin mutant mice
S/A_1283_: Reelin with the serine to alanine mutation at position 1283
VLDLR: very low-density lipoprotein receptor

