## Supplementary Figures and Methods for "Serine 1283 in extracellular matrix glycoprotein Reelin is crucial for Reelin’s function in brain development"

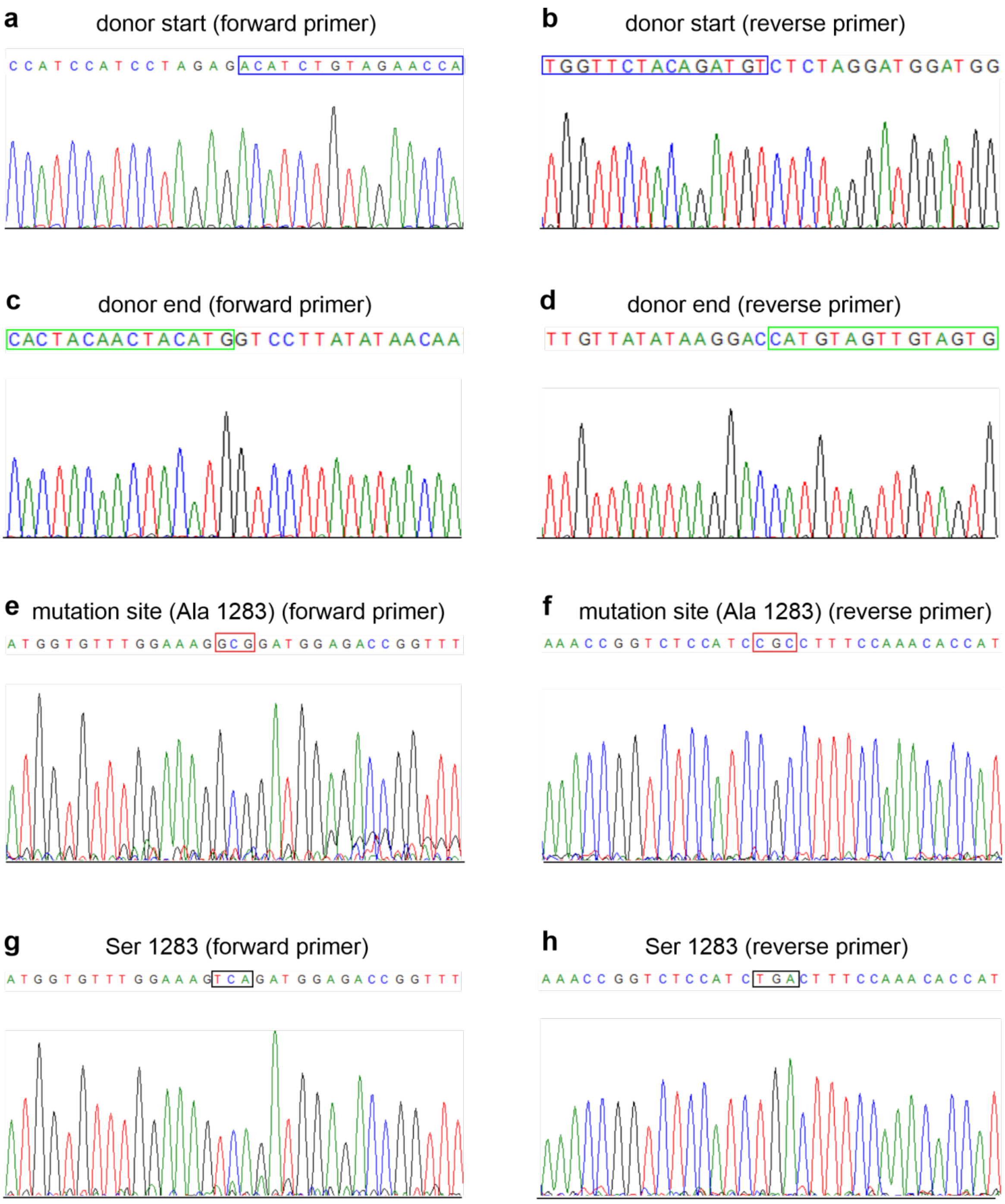


**Fig. S1. Gene sequencing.** Genomic DNA isolated from *rln*$^{+/+}$ and *rln*$^{A/A}$ brains and primers covering the donor DNA sequence and sequences 341 bp upstream and 483 bp downstream thereof were used. (**a-d**) Representative chromatograms from the sequenced part of the Reelin gene of *rln*$^{A/A}$ mice using forward and reverse primers. Sequenced stretches from the start (**a**, **b**) and end (**c**, **d**) of the donor sequence and flanking sequences as well as from the mutation site (**e**, **f**) are shown. (**g**, **h**) Chromatograms and sequence stretches of the mutation site from sequencing of genomic DNA from a *rln*$^{+/+}$ mouse are shown.

**Serine 1283 in extracellular matrix glycoprotein Reelin is crucial for Reelin´s function in brain development**

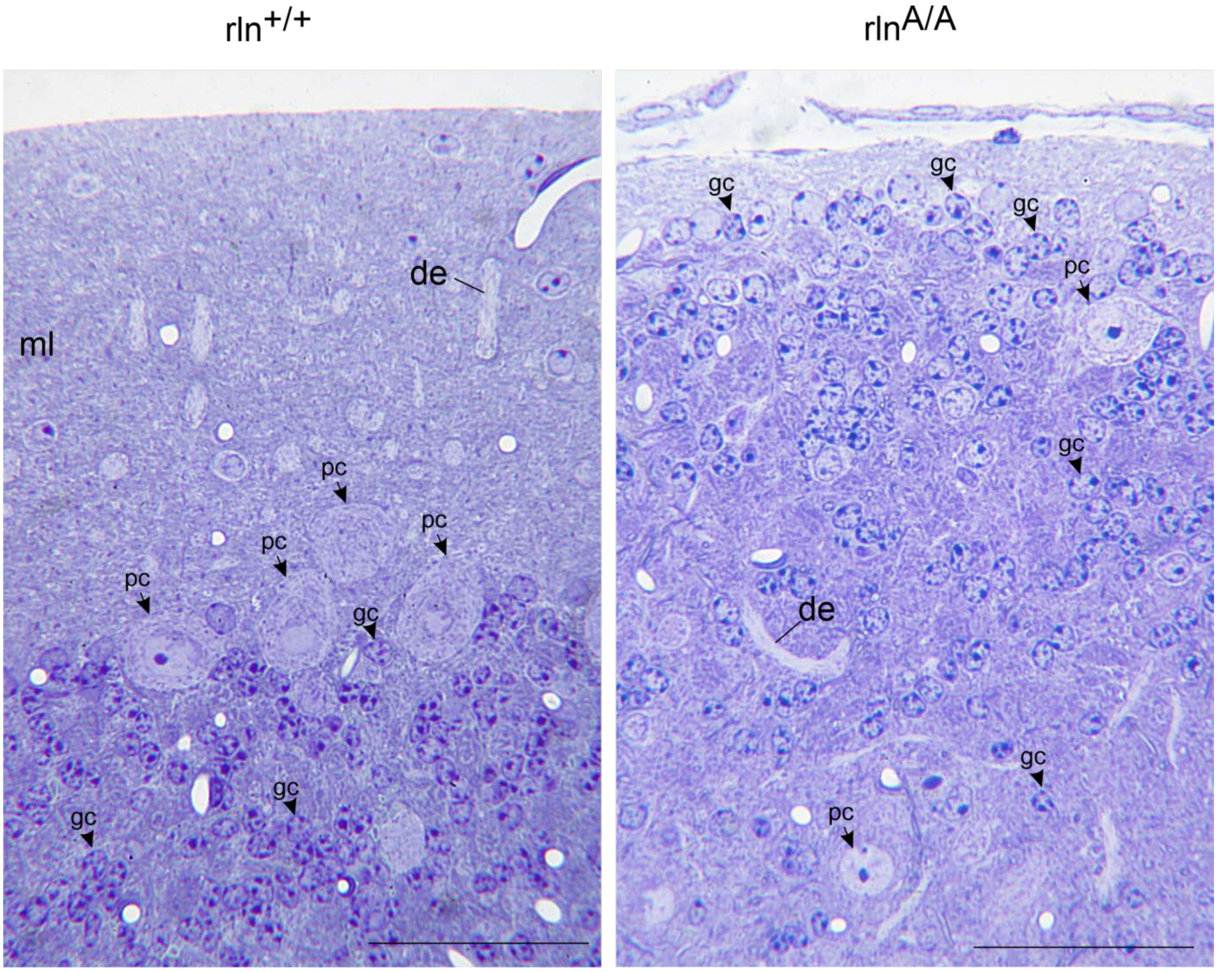


**Fig. S2. Toluidine blue staining of cerebella from one-month-old *rln*$^{+/+}$ and *rln*$^{+/A}$ mice**. In the *rln*$^{A/A}$ cerebellum misplaced Purkinje cells (arrows; pc) and granule cells (arrowheads; gc) were observed. de, dendrite of the Purkinje cell; ml, molecular layer; scale bars: 50 µm.

### Serine 1283 in extracellular matrix glycoprotein Reelin is crucial for Reelin´s function in brain development

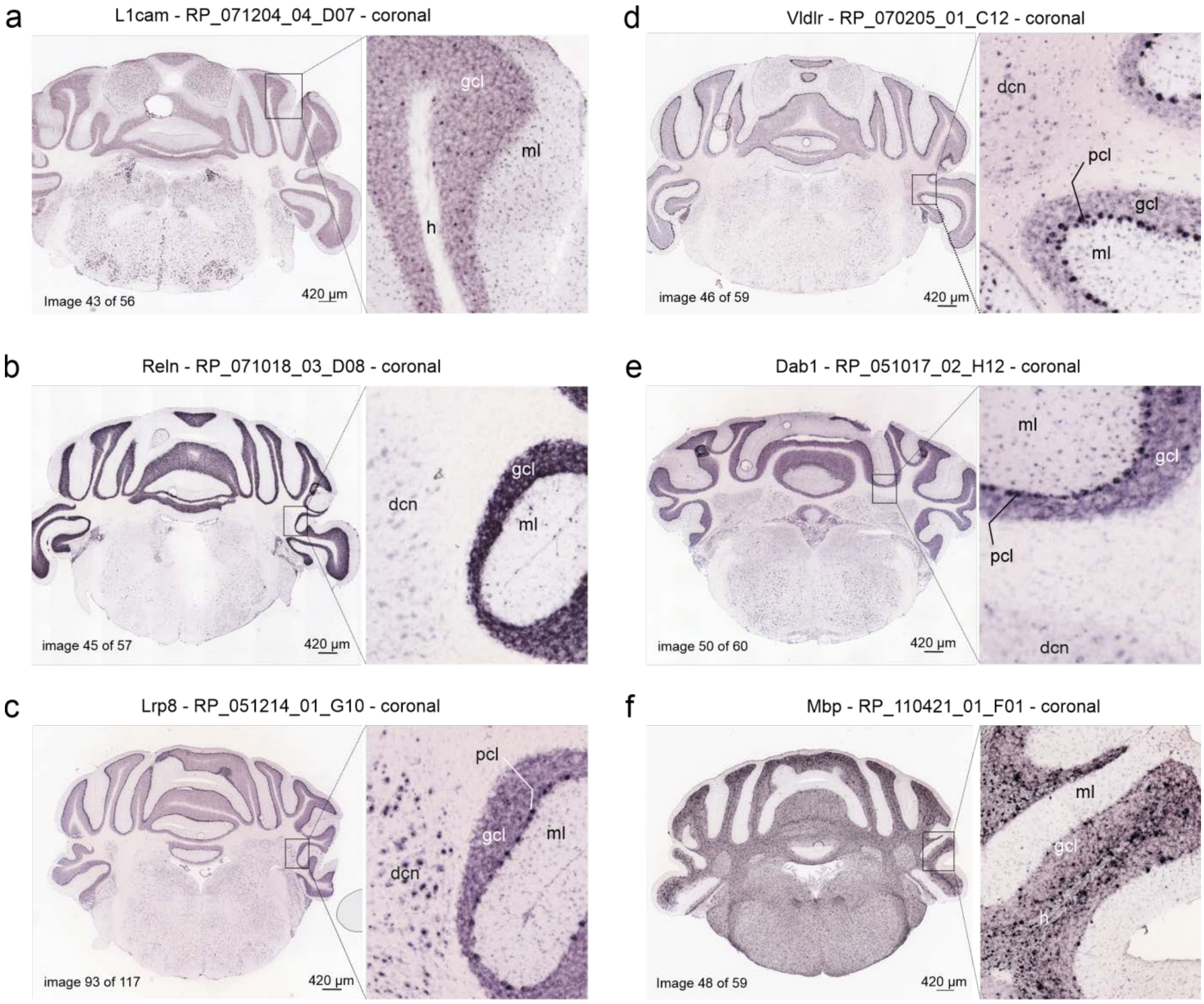


**Fig. S3. mRNA expression of L1, Reelin, ApoER2/Lrp8, VDLR, Dab1 and MBP in cerebellar granule cells**. Shown are *in situ* hybridization images of coronal sections from the Allen Brain Atlas (© 2004 Allen Institute for Brain Science. Allen Mouse Brain Atlas. Available from: mouse.brain-map.org) for L1 **(a)**, Reelin **(b)**, ApoER2/Lrp8 **(c)**, VDLR **(d)**, Dab1 **(e)** and MBP **(f)**. At a higher magnification the images from the cerebellum of an adult mouse (postnatal day 57) granule cells appear to be positive for Reelin, Dab1, ApoER2, VDLR, MBP or L1. dcn, deep cerebellar nuclei; gcl, granule cell layer; h, hilar region; ml, molecular layer; pcl, Purkinje cell layer.

**Supplementary Methods**

#### Generation of gene-edited mice, Southern blot analysis, genomic sequencing and genotyping

The wild-type Reelin sequence on mouse chromosome 5 (NC_000071.6; Gene ID: 19699) was used for the design and screening of suitable zinc finger nuclease pairs (Sigma-Aldrich), which bind at position 21,998,036-21,998,022 (ZFN1) and 21,998,061-21,998,043 (ZFN2) and close to the TCA codon coding for serine 1283 at position 21,997,954-21,997,952. In the mutated sequence, the serine codon TCA was replaced by GCG (coding for alanine) generating thereby a restriction site for AciI. After confirmation of the zinc finger nuclease specificity, the zinc finger nuclease plasmids were linearized, purified by phenol-chloroform extraction, precipitated with ethanol, transcribed, tailed according to the manufacturer's instructions (Epicentre, Madison, WI, USA; MMA60710 and PAP5104H), and purified using the MEGA clear Kit (Ambion, ThermoFisher Scientific, Darmstadt, Germany). The zinc finger nuclease transcripts and a 1,600 bp donor sequence (21,997,240-21,998,839) carrying the exchanged codon GCG in a pUC-vector (Genewiz, Leipzig, Germany) were injected under constant pressure into the pronuclei of one-cell-stage embryos derived from C57BL6/JxCBA mice and implanted into foster mothers according to standard procedures[1]. The correct insertion of the mutation into the genomic DNA was confirmed by Southern blot analysis and sequencing using genomic DNA and primers covering the donor sequence and sequences 341 bp upstream and 483 bp downstream thereof.
For genotyping, mouse tail biopsies were taken and immediately processed using the Phire Animal Tissue Direct PCR Kit (ThermoFisher Scientific) according to the manufacturer's instructions. Primers P1 (5'-CTT ATG TTT TTG TGT TTT CAG TGT-3'; 21,998,152-21,998,129) and P2 (5'- GTT CAA AAC ATC ACT CTC AGA TCA -3'; 21,997,623-21,997,646) were used for amplification of a 530 bp region according to the following program: 98°C for 5 min; 35 cycles: 98°C for 10 s, 57°C for 45 s and 72°C for 20 s; 72°C for 7 min; 4°C until further use. The amplification product was digested with AciI (New England Biolabs, Frankfurt am Main, Germany) in CutSmart buffer (New England Biolabs) at 37°C for 2 h. The 530 bp amplicon was cleaved into 199 bp and 331 bp products visualized on a 2.5% agarose gel containing ethidium bromide.
For Southern blot analysis, genomic DNA was isolated from frozen brain tissue using the DNA Isolation Reagent for Genomic DNA (A3418; AppliChem, Darmstadt, Germany), digested with EcoRI, separated on a 0.8% agarose gel (Invitrogen), and transferred to Amersham Hybond-N+ nylon membrane (Cytiva Europe, Freiburg, Germany) (Herrmann et al., 1986). The donor plasmid and the primers P1 and P2 were used for PCR to amplify a 530 bp probe. The probe was run on a 2% agarose gel, purified using NucleoSpin® Gel and PCR Clean Up kit (Macherey-Nagel, Düren, Germany), labeled with $^{32}$P-dATP (Hartmann Analytic, Braunschweig, Germany) using the Amersham Megaprime DNA Labelling System (Cytiva), and purified using a G50 spin column. Hybridization was performed overnight at 65°C (Church and Gilbert, 1984). After washing three times at 65°C, the membrane was exposed to a phosphoimager-plate and analyzed with a phosphoimager (Fujix Bas 2000, Fuji Photo Film, Tokyo, Japan).
For genomic sequencing, genomic DNA purified from $rln^{+/+}$ and $rln^{A/A}$ mouse brains was used to generate an amplicon of 2424 bp using the following primers: PCR fw (GTC ATC AGA CTT GGC GAC AGA) and PCR rev (CCT ATC ATC AAC CAG CGA GC). The amplicon was then sequenced with the following primers: seq fw1 (TGA CCA TTG TAT GAC CAA CAT GAC), seq fw2 (AAA CAT GAA TGG TGT ATG GCC CAC), seq fw3 (TTA GAA TTC CCT CCC TGT AAC TTG), seq rev1 (TCA GGA TTT AAT GTT TAC

CCT GCC), seq rev2 (TTC GCC ATC CTA AGC CTG TG), and seq rev3 (TCT TCA GGA AGC CAG TCA TTC).
For whole genome sequencing, 1 µg of genomic DNA purified from *rln*$^{+/+}$ and *rln*$^{A/A}$ mouse brains was sent to commercial sequencing (Novogene, Cambridge, UK) to confirm the correct insertion of the Reelin mutation $S/A_{1283}$ and the correct positioning of the Reelin sequence within the Reelin gene by superposing the *rln*$^{A/A}$ DNA sequence against the *rln*$^{+/+}$ sequence.

**Pencil grab test and assessment of general locomotion**

For locomotion assessment, one-month-old female *rln*$^{+/+}$ and *rln*$^{A/A}$ mice were used. For the pencil grab test, a wooden pencil of 15 cm length and diameter of 7 mm was mounted vertically (tip of the pencil facing upwards). Mice were held on their tails with the head facing downwards, at a distance of 1 cm from the pencil and a depth position of the head of 5 cm from the tip of the pencil. The *rln*$^{+/+}$ mice grabbed the pencil with their forepaws and thereby clapped their hindpaws together. The *rln*$^{A/A}$ mice were analyzed for deviation in their hind limbs coordination in respect to the *rln*$^{+/+}$ behavioral reference.
One-month-old female *rln*$^{+/+}$ and *rln*$^{A/A}$ mice were placed pairwise in empty cages and observed for 30 min. The stance and posture of the body of the *rln*$^{+/+}$ mice were used as a reference for the determination of deviation in body stance and posture in *rln*$^{A/A}$ mice. The *rln*$^{+/+}$ mice explored the cage by "rearing", thereby standing upright on their hindpaws and holding their forepaws on the wall of the cage. This behavior was the reference for the assessment of the explorative behavior of *rln*$^{A/A}$ littermates.
For assessment of body coordination, one-month-old female mice of both genotypes were placed on the barred cage lid and observed for 30 min. The *rln*$^{+/+}$ mice could balance on the bars and on the edges of the cage, and when placed at the lid swale manage to climb to the plateau. This behavior was used as a reference for assessment of deviating coordination in *rln*$^{A/A}$ littermates.

**Isolation of RNA, reverse transcription and quantitative real-time PCR**

Total RNA was isolated from brains using the TRI reagent (Sigma-Aldrich) and RNAeasy Plus Kit (Qiagen, Hilden, Germany) following the manufacturer's instructions. For reverse transcription, 0.5-5 µg total RNA, 0.1 pM oligoT18 primer and 200 U M-MLV reverse transcriptase (Sigma-Aldrich) were used. Quantitative real-time PCR was performed in triplicates using the 7900HT Fast Real-Time PCR System (ThermoFisher Scientific), the qPCR kit SYBR® Green I, ROX (Eurogentec, Cologne, Germany), reverse transcribed RNA (1:10 diluted), and 0.1 pM primers for determination of the mRNA levels of Reelin (5'- AAC TTC TAT GAG AAG CCA GCT TTC -3'; 5'- AAC TGC AGC ACA TAT CCA GGT TTC -3') or of the reference genes actin (5´- TCC TGT GGC ATC CAT GAA ACT -3´, 5´-TTC TGC ATC CTG TCA GCA ATG-3´), tubulin (5´-CGC ACG ACA TCT AGG ACT GA-3´, 5´-TGA GGC CTC CTC TCA CAA GT-3´), and hypoxanthine phosphoribosyltransferase 1 (5´-GTT CTT TGC TGA CCT GCT GGA-3´, 5´-TCC CCC GTT GAC TGA TCA TT-3´). The SDS 2.4 software was used for analysis of data from quantitative real-time PCR. The mRNA levels of Reelin relative to the mRNA levels of the reference genes were calculated. Data of relative gene expression at the mRNA level were analyzed by two-way analysis of variance (ANOVA) followed by Bonferroni post-hoc test.

**Site-directed mutagenesis**

For the serine-to-alanine exchange at position 1283 in Reelin, a sequence of 5,000 bp in the region between SnaBI and AgeI restriction sites of the full-length Reelin vector was amplified with the HiFi amplification kit (Clontech, Takara Bio Europe, Saint-Germain-en-Laye,

France) using the forward primer 5’-AAC CCC ACC TAC TAC GTA CCG GGA CAG GAA TAC-3’ and the mutated reverse primer 5’-GT TAC TGC AAA CCG GTC TCC ATC CGC CTT TCC AAA CAC CAT GGC TG-3’ (underlined triplet indicates the mutated region). The full-length Reelin vector was digested with enzymes SnaBI and AgeI, and the mutated 5,000 bp amplicon was fused with the appropriate vector using the In Fusion Cloning Kit (Clontech) according to the manufacturer’s instructions.

#### *In utero* electroporation

Timed pregnant mice at E13.5 were anaesthetized by inhalation anesthesia with 4% isoflurane vaporized in an $O_2$-volumetric flow of 0.8 l/min. During surgery, the isoflurane concentration was lowered to 1.5-2%. The abdominal cavity of the pregnant mice was opened, and the uterine horns bearing the embryos were exposed.

For *in utero* electroporation and rescue experiments using *rln*$^{A/A}$ embryos, 2 µg of plasmid coding for N-R6 and 2 µg of *CAG-IRES-GFP* vector were pre-mixed with 0.01% Fast Green and unilaterally injected into the ventricle of embryonic brains. Embryos were electroporated with five electrical square unipolar pulses (amplitude of 30 V, duration of 50 ms, intervals of 950 ms) powered by a BTX electroporation apparatus (model BTX ECM 830; Harvard Apparatus, Holliston, MA, USA). The uterine horns were placed back into the abdominal cavity and mice were allowed to survive for 60 h. The pregnant mice carrying the electroporated embryos were then sacrificed and embryos were isolated in ice-cold phosphate buffered saline, pH 7.3 (PBS) and killed by decapitation. Tail biopsies from the embryos were taken for genotyping. Brains electroporated to express GFP were collected under a fluorescence microscope and fixed overnight in 4% w/v formaldehyde in 0.1 M sodium cacodylate buffer, pH 7.4 (Sigma-Aldrich) at 4°C, followed by an overnight immersion at 4°C in a solution of 15% w/v sucrose in 0.1 M sodium cacodylate buffer, pH 7.4.

For standard fluorescence microscopy, electroporated brains were embedded in 3% agarose and sectioned on a vibratome (Leica VT1000 S; Leica Instruments, Wetzlar, Germany) at a thickness of 80 µm and immersed in blocking solution (5% normal goat serum and 0.2% Triton-X 100 in 0.1 M PBS), washed, and incubated with Reelin antibody G10 (dilution 1:500) at 4°C overnight. The tissue sections were washed in PBS, incubated in appropriate secondary antibodies (dilution 1:250) at 4°C overnight, counterstained with DAPI to label nuclei, mounted in Dako Anti-fade Fluorescence Mounting Medium (Dako, Agilent, Santa Clara, CA, USA), and subjected to fluorescence microscopy.

#### Tissue preparation, immunohistology and imaging

Brain slices of 25 µm thickness were prepared as described[2]. Sections, stored at -20°C, were air-dried for 30 min at 37°C and immersed for antigen demasking in 0.01 M sodium citrate solution (pH 9.0, adjusted with 0.01 M NaOH), preheated to 80°C in a water bath for 30 min. Following cooling to room temperature, blocking of non-specific binding sites was performed using PBS supplemented with 0.2% Triton X-100 (VWR, Darmstadt, Germany), 0.02% sodium azide and 5% donkey serum (ThermoFisher Scientific) at room temperature for 1 h. For paraffin sectioning, fixed brains were incubated in 70% ethanol at 4°C overnight. Brains were dehydrated in ascending grades of ethanol and cleared in pure xylene. Following immersion in liquid paraffin (Sigma-Aldrich) at 60°C for 24 h, samples were embedded in paraffin at ambient room temperature. Sections of 10 µm were cut on a microtome (Leica VT1000 S) and collected on glass slides. Sections were deparaffinized in pure xylene several times, rehydrated in descending grades of ethanol and subjected to immunohistochemistry. Immunohistochemistry was performed with the following primary antibodies, which were diluted in blocking solution (PBS containing 10% horse serum and 0.01% Triton X-100): Reelin antibody G10 (1:500), rabbit polyclonal MBP antibody (1:500), NeuN antibody

(1:1,000), and antibodies directed against Brn2 (1:250), calbindin (1:1,000) or Cux1 (1:250). Secondary donkey antibodies conjugated with Cy3 and Cy2 were used at a dilution of 1:500 in PBS. Sections were mounted in anti-fading medium Fluoromount containing DAPI (Sigma-Aldrich) and stored in the dark at 4°C.

Images of immunostained brain sections were taken on a confocal fluorescence microscope (FluoView FV1000, Olympus, Hamburg, Germany) or a Keyence Fluorescence Microscope (BZ-9000, Keyence, Neu-Isenburg, Germany) and were processed with ImageJ (ImageJ, RRID:SCR_003070).

For semithin sectioning and toluidine blue staining, 4% formaldehyde-fixed cerebella from one-month-old female *rln*$^{+/+}$ and *rln*$^{A/A}$ animals were osmicated with 2% aqueous $OsO_4$ solution for 2 hours at room temperature, dehydrated in ascending grades of ethanol and pure propylene and then embedded in Araldite resin. Semithin sections of 0.75 µm thickness were prepared and stained with 1% toluidine blue in PBS. Sections were subjected to light microscopy using a 40× objective and the Leica Light Microscope DM500 equipped with a Canon EOS 70D camera.

**Counting of Cajal-Retzius (CR) cells in *rln*$^{+/+}$*-CXCR4-GFP* and *rln*$^{A/A}$*-CXCR4-GFP* mouse cerebellar cortices**

Mounted sagittal brain slices from 24-day-old female *rln*$^{+/+}$*-CXCR4-GFP* and *rln*$^{A/A}$*-CXCR4-GFP* littermates were subjected to fluorescence microscopy with focus on the marginal zone of the somatosensory cortex in *rln*$^{+/+}$*-CXCR4-GFP* mice and on the corresponding analogous region in *rln*$^{A/A}$*-CXCR4-GFP* mice. The marginal zone in the wild-type and its analog in the mutant mice was divided in four to five segments, each segment with a depth of 70 µm (from the pial surface) and width of 50 µm. Each segment was divided into seven bins of 10 µm depth. The CR cells were counted in each bin and the CR cell distribution across the seven bins of each segment was assessed. The difference between values of the means + SEM (sample size of n = 18 for each group; 3 sections from 6 animals per genotype) among both genotypes were tested with two-way ANOVA for multiple comparisons and Sidak's multiple comparisons test. Moreover, for a sample size of n = 18 for each genotype the number of CR cells was estimated within at least 5 legs of 50 µm of the 70 µm-deep superficial layer of the CR cells. The difference between values of the means + SEM of the number of CR cells per 50 µm leg among both genotypes was tested with the two-tailed t-test.

**Nissl staining**

Paraffin-embedded brains were cut sagittally into 10 µm-thick sections. Sections were incubated in 100% xylene for 10 min and rehydrated in 100%, 90%, 70% ethanol for 5 min each at room temperature. Sections were stained with cresyl violet according to standard Nissl protocols, mounted on coverslips and imaged on the Keyence Microscope BZ-9000.

**Preparation of brain homogenates, SDS-gel electrophoresis, silver staining, immunoblot analysis and immunoprecipitation**

Mice younger than postnatal day 14 were sacrificed by decapitation. Older mice were anaesthetized with a gas mixture of 80% $CO_2$ and 20% $O_2$ prior to cervical dislocation. Brains were dissected in ice-cold PBS. Brain areas of interest were carefully excised and homogenized in RIPA buffer consisting of 20 mM Tris-HCl at pH 7.5, 150 mM NaCl, 1 mM $Na_2EDTA$, 1 mM EGTA, 1% Nonidet P-40, 1% sodium deoxycholate, 2.5 mM sodium pyrophosphate, 1 mM β-glycerophosphate, 1 mM $Na_3VO_4$, and 1× cocktail of protease inhibitors (Sigma-Aldrich) at 4°C. After centrifugation at 20,000 × g and 4°C for 10 min, supernatants were collected and equal amounts of total protein from the supernatants were subjected to SDS-gel electrophoresis and immunoblot analysis as described[4] using Reelin antibody G10 (diluted 1:1,000), L1 antibody C-2 (diluted 1:1,000), goat polyclonal MBP antibody (diluted 1:1,000), phospho-Ser antibody (1:1,000) and GAPDH antibody (diluted 1:2,000). Secondary antibodies coupled to horseradish peroxidase were used at a dilution of 1:10,000.

For immunoprecipitation, cell culture supernatants of cultured cerebellar neurons were incubated overnight with Reelin antibody G10 followed by overnight incubation with Protein A/G beads (Santa Cruz Biotechnologies). After washing three times with PBS, the beads were boiled 5 min at 95°C. The supernatant was then subjected to SDS-gel electrophoresis and immunoblot analysis.

For silver staining, samples were run on SDS-gels and stained with Pierce™ Silver Stain for mass spectrometry (ThermoFisher Scientific).

#### Purification of wild-type and $S/A_{1283}$-mutated Reelin

Wild-type Reelin and Reelin with the serine to alanine mutation at position 1283 ($S/A_{1283}$) were purified from transfected HEK cell culture supernatants[3] using Heparin High-Trap Sepharose columns (GE Healthcare Life Sciences, Freiburg, Germany) following the manufacturer's instructions (see also Lutz et al.[4]). Supernatants from mock-transfected HEK cells served as controls. Elution was performed in two steps using 140 mM and 400 mM NaCl solution. The eluates containing Reelin were dialyzed using 100 kDa cut-off columns (Amicon/Merck). The purified protein fractions were subjected to electrophoresis in SDS-polyacrylamide gels that were either silver-stained or used for immunoblot analysis with the G10 antibody to measure protein yields and protein purity (see Lutz et al.[4], Supplementary Figure S4).

#### Dissociated cell cultures, *$rln^{A/A}$-POMC-GFP* explants and imaging of migrating *POMC-GFP*-positive cerebellar granule cells

Isolation and culturing of murine cortical neurons, and murine cerebellar neurons and explants have been described[5-7]. In brief, murine cerebral cortices were dissected from embryonic C57BL/6 brains at the desired developmental age and incubated in 0.025% trypsin (Sigma-Aldrich) in HBSS at 37°C for 30 min. The cortices were then incubated in HBSS containing 1% BSA (Sigma-Aldrich) and 1% w/v trypsin inhibitor (T-6522, Sigma-Aldrich) at 37°C for 5 min. After washing in HBSS, the tissue was mechanically dissociated and the dissociated neurons were cultured in Neurobasal medium (Invitrogen) supplemented with 1% B-27 (Invitrogen), 2 mM L-glutamine (Invitrogen), 100 units/ml penicillin (Invitrogen), and 100 µg/ml streptomycin (Invitrogen) at a density of $2 \times 10^6$ cells per well of a 6-well plate coated with poly-L-lysine (Sigma-Aldrich).

Cerebellar neurons were prepared from cerebella of 6- to 8-day-old mice of the desired genotype. Dissected cerebella were incubated with 10 mg/ml trypsin and 0.5 mg/ml DNase I (Sigma-Aldrich) in HBSS for 15 min at 37°C, washed with HBSS, mechanically dissociated and centrifuged at 100 g for 15 min. Dissociated cells were diluted in culture medium containing Neurobasal A medium (Invitrogen) supplemented with 2 mM L-glutamine (Invitrogen), 4 nM L-thyroxine (Sigma-Aldrich), 1 mg/ml BSA (Sigma-Aldrich), 12.5 µg/ml insulin (Sigma-Aldrich), 30 nM sodium selenite (Sigma-Aldrich), 100 µg/ml transferrin (Merck Biosciences), 0.1 mg/ml streptomycin, 10 U/ml penicillin (Invitrogen), and seeded at a density of $3 \times 10^6$ cells per well of a 6-well plate (immunoblot analysis) or $2.5 \times 10^5$ cells per well of a 48-well plate (cell death analysis) coated with poly-L-lysine (Sigma-Aldrich).

For treatment with CK2 inhibitors, cerebellar neurons were maintained in serum-free medium overnight. Neurons were then incubated for 2 h in fresh serum-free medium containing 25 µM TBB and 0.025 % DMSO, 500 nM K252b and 0.025 % DMSO or only 0.025 % DMSO. Cell

culture supernatants were collected and centrifuged at 10,000 g and 4°C, while cells were resuspended in 150 µl RIPA buffer. The cell-free supernatant and cell lysates were subjected to immunoblot analysis. Cell survival was determined using neutral red uptake assay after treatment of cerebellar neurons with 25 µM TBB and 500 nM K252b for 30 h.
To characterize Reelin secretion in cell culture, cerebral cortex neurons were maintained overnight, and cell culture supernatants were collected, centrifuged at 10,000 g and 4°C, and subjected either to protein precipitation using methanol-chloroform[8] or to immunoprecipitation using Dynabeads Protein G (ThermoFisher Scientific) and Reelin antibody G10. Precipitated proteins, immunopurified Reelin and cells lysed in RIPA buffer were resuspended in SDS-sample buffer and subjected to immunoblot analysis.
Cerebellar explants were prepared from *rln*$^{+/+}$*-POMC-GFP* and *rln*$^{+/+}$*-POMC-GFP* neonates to measure neuronal migration which was determined from at least 12 explants per treatment and experiment. Briefly, cerebella were forced through Nitrex nets with pore sizes of 300, 200 and 100 µm. The cerebellar pieces (explants) were incubated with 5 ng of purified wild-type or $S/A_{1283}$-mutated Reelin and then seeded onto Matrigel-coated coverslips, and incubated at 37°C in a 5% $CO_2$ containing atmosphere for 3 h before real-time microscopy. Real-time microscopy and quantitative analyses were performed with an Improvision spinning disk microscope (Carl Zeiss/Perkin Elmer, Waltham, MA, USA) equipped with a 488-nm scanning laser and an incubation chamber containing an inner chamber with saturated humidity and 5% $CO_2$ atmosphere at 37°C. For imaging, a 20× objective lens (numerical aperture: 0.5; immersion medium: air) was used. Time-lapse images were collected every 20-40 min for 8 h. Photobleaching during imaging was prevented by minimizing the laser power. Initial image processing was carried out with Volocity 6.1.1 (Perkin Elmer). TIFF-series at time rates of 3 frames/sec were integrated and video-edited.

#### *In vitro* phosphorylation assay using CK2 and purified wild-type and $S/A_{1283}$-mutated Reelin

For phosphorylation, 1 µg wild-type Reelin or $S/A_{1283}$-mutated Reelin was incubated for 1 h at 30°C in 100 µl containing 100 µM ATP, 10 µCi ɣ-$P^{32}$-ATP (Hartmann Analytics, Braunschweig, Germany), 500 U CK2 (New England Biolabs), and 1x NEBuffer for Protein Kinases (New England Biolabs). The samples were then subjected to protein precipitation using methanol-chloroform[8], and the precipitates were resuspended in SDS sample buffer and then subjected to SDS-PAGE followed by blotting onto nitrocellulose membrane. Radiolabeled proteins were visualized by a Phosphor-Imager (Molecular Dynamics, Sunnyvale, USA).

#### ELISA

For ELISA, 250 ng MBP or 2.5 µg heparin-purified wild-type or mutated Reelin were substrate-coated in 384-well microtiter plates with high binding surface (Corning, Wiesbaden, Germany) at 4°C overnight. After washing with PBS, blocking with 1% w/v bovine serum albumin (essentially fatty acid free; Sigma-Aldrich) in PBS at room temperature for 1 h, and washing with PBS, different concentrations of purified wild-type Reelin (10, 25, 50, 100, 200 µg/ml) or MBP (1, 2.5, 5, 10, 20 µg/ml) were added to the substrate-coats for incubation at room temperature for 2 h. After washing with PBST (PBS with 0.05% Tween 20), mouse monoclonal Reelin antibody G10 (1:100) and goat polyclonal MBP antibody (1:100) followed by an anti-mouse and anti-goat HRP-coupled secondary antibody (1:2,000; Dianova) as well as O-phenylenediamine dihydrochloride (ThermoFisher Scientific, Darmstadt, Germany) as horseradish peroxidase (HRP) substrate were used for detection of Reelin and MBP. The reaction was terminated by addition of 2.5 M sulphuric acid. Absorbance was measured at 492 nm with the ELISA reader (BioTek, Bad Friedrichshall, Germany).
